# Focusing on human haplotype diversity in numerous individual genomes demonstrates an evolutional feature of each locus

**DOI:** 10.1101/2020.03.28.012914

**Authors:** Makoto K. Shimada, Tsunetoshi Nishida

**Author notes:** Author for Correspondence: Makoto K. Shimada, Department of Gene Expression Mechanism, Institute for Comprehensive Medical Science, Fujita Health University, Aichi 470-1192, Japan.

## Abstract

The application of current genome-wide sequencing techniques on human populations helps elucidate the considerable gene flow among genus *Homo*, which includes modern and archaic humans. Gene flow among current human populations has been studied using frequencies of single nucleotide polymorphisms. Unlike single nucleotide polymorphism frequency data, haplotype data are suitable for identifying and tracing rare evolutionary events. Haplotype data can also conveniently detect genomic location and estimate molecular function that may be a target of selection. We analyzed eight loci of the human genome using the same procedure for each locus to infer human haplotype diversity and reevaluate past explanations of the evolutionary mechanisms that affected these loci. These loci have been recognized by separate studies because of their unusual gene genealogy and geographic distributions that are inconsistent with the recent out-of-Africa model. For each locus, we constructed genealogies for haplotypes using sequence data of the 1000 Genomes Project. Then, we performed S* analysis to estimate distinct gene flow events other than out-of-Africa events. Furthermore, we also estimated unevenness of selective pressure between haplotypes by Extended Haplotype Homozygosity analysis. Based on the patterns of results obtained by this combination of analyses, we classified the examined loci without using a specific population model. This simple method helped clarify evolutionary events for each locus, including rare evolutionary events such as introgression, incomplete lineage sorting, selection, and haplotype recombination that may be hard to discriminate from each other.

## Introduction

Recent advancements in ancient genomics have provided unprecedented insights into ancient population dynamics, which include migration and bi-directional gene flow of archaic human groups, such as Neandertals and Asian *Homo erectus* (Green et al. 2010; Reich et al. 2011; Meyer et al. 2012; Fu et al. 2014; Prufer et al. 2014; Kuhlwilm et al. 2016). This information revealed that genomes of modern human populations contained various genomic fragments that originated from archaic humans, and some studies suggested that introgressed genomic fragments are adaptive in certain environments (Sankararaman et al. 2014; Racimo et al. 2016; Sankararaman et al. 2016; Simonti et al. 2016; Dannemann and Kelso 2017; Enard and Petrov 2018)(rev., Dannemann and Racimo 2018).

Furthermore, recently developed sequencing technology has changed data in format and quantity, which has prompted innovation of data analysis. Data used in human genome evolution can be classified into two types: frequency and sequence data.

Single nucleotide polymorphism (SNP) frequency in current populations has been mainly processed to analyze genomic relationships among human populations in most human genomics studies (Harris and Michael 2017). Research using SNP frequency data can address genetic similarities among populations, population structure, effective population sizes, gene flow, and selection; additionally, disease-causing variants have been suggested by genome-wide association studies. Because frequency data are advantageous in quantitative analyses, they have also been applied to estimate the extent of gene flow among populations (e.g., Mallick et al. 2016; Mondal et al. 2016; Jinam et al. 2017; Lipson and Reich 2017).

Although SNP data were treated as independent from each other, haplotype sequence data are a collective body of neighboring SNPs that share common evolutionary history and molecular function. Accordingly, haplotype data comparatively easily connect to information about sequence motifs and genomic position. If simple, established methods that use haplotype data are available in population genomics, researchers can estimate molecular function of genomic regions that were suggested to be introgressed from other populations and selected for in the introgressed population.

Sequencing technology advancements have also facilitated genome-wide studies that effectively reveal past demographic process without bias in choosing target locus. Differing from demographic events, however, introgression events leave a signal, and fragmented sequences can be detected in the form of a patchwork or mosaic through recombination and drift at each locus. Consequently, the accumulated locus-oriented studies are indispensable for characterizing and comprehending gene flow and allele maintenance mechanisms (Mendez et al. 2012a). Additionally, even before the first genome-wide sequencing of archaic humans by Green et al. (2010), some studies on modern humans claimed that unusual haplotypes were inconsistent with the recent out-of-Africa (OOA) model based on their gene genealogy and geographic distribution pattern (Zietkiewicz et al. 2003; Garrigan et al. 2005a; Garrigan et al. 2005b; Hardy et al. 2005; Stefansson et al. 2005; Shimada et al. 2007). Such diversified haplotypes within modern human population genomes have been studied separately (Evans et al. 2006; Shimada et al. 2007; Cox et al. 2008; Donnelly et al. 2010; Yotova et al. 2011; Mendez et al. 2012b; Ding et al. 2013; Mendez et al. 2013). Recently, a lot of individual human genomes have been sequenced, including those of archaic humans, which has allowed researchers to evaluate the origin of genome-scale variation using a unified method. However, the following problems hamper the use of haplotype data from a large number of individual genomes. First, there is no practical definition of a genomic region of interest (locus) from the massive amount of genome-wide data. A sequence-based analysis, such as gene genealogy, uses the locus as a specifically defined unit of sequence alignment. To ensure accuracy, a longer genomic region with a larger data set that shares common evolutionary history should be selected as a locus to be analyzed. Long sequences without recombination hotspots were preferred in previous studies to obtain a better estimate of TMRCA in population genetic analysis (Cox et al. 2008). The available genome-wide data sets of individuals from multiple populations contain recombined haplotypes that have independently recombined in various genomic positions. Furthermore, a large amount of longer sequence data sets may be more frequently influenced by factors such as inversion, gene duplication, copy number variation, and selection. Accordingly, there should be focus on developing a definition of a locus. Second, there is no method to distinguish between introgression and incomplete lineage sorting (ILS) of ancestral polymorphisms. Genomes are thought to contain both genomic fragments that were derived from introgression and retained via ILS (Wang et al. 2018). Furthermore, different gene genealogical patterns are expected depending on order of coalescence and population division (Joly et al. 2009). Accordingly, discrimination between them is impossible by gene genealogy alone. As larger data sets are used, such difficulties are expected to be encountered with a higher frequency. Previous studies have suggested that a simple dichotomic framework is not sufficient to judge ILS or introgression in Eurasia after OOA (e.g., Shimada et al. 2007; Campbell and Tishkoff 2010; Lipson and Reich 2017; Povysil and Hochreiter 2017). A specific model-based verification method cannot always be applied for all evolutionary events that have been experienced in human populations. Consequently, a simple, model-free method with fewer assumptions is needed to focus on questions regarding the development of population genomics with a large amount of individual genome-wide data.

The purpose of this study was to provide various examples of human genome diversity using a single combination of haplotype-based methods. This will help: 1) define a locus in genome-wide sequence data from a massive amount of individuals, and 2) compare and reevaluate differences in haplotype variation among genomic segments that reflect evolutionary history. Therefore, we focused on eight loci that have been noted to have unusual gene genealogy and/or geographic distributions inconsistent with the OOA model (Table 1). Using a public catalog of human variation, the 1000 Genomes Project, as a common data set for these eight loci, we demonstrated haplotype genealogy with estimation of haplotype-specific selection and introgression from known and unknown archaic humans. Using S* analysis, we estimated introgression from archaic hominins found in the 1000 Genome Project samples for these eight loci. We also evaluated unevenness of selective pressure between the most diverged haplotypes and other haplotypes across the examined loci using Extended Haplotype Homozygosity (EHH) analysis. These strategies demonstrated that genomes of human populations contain various backgrounds, and this approach represents a possible method to distinguish introgression from ILS.

**Table 1.**
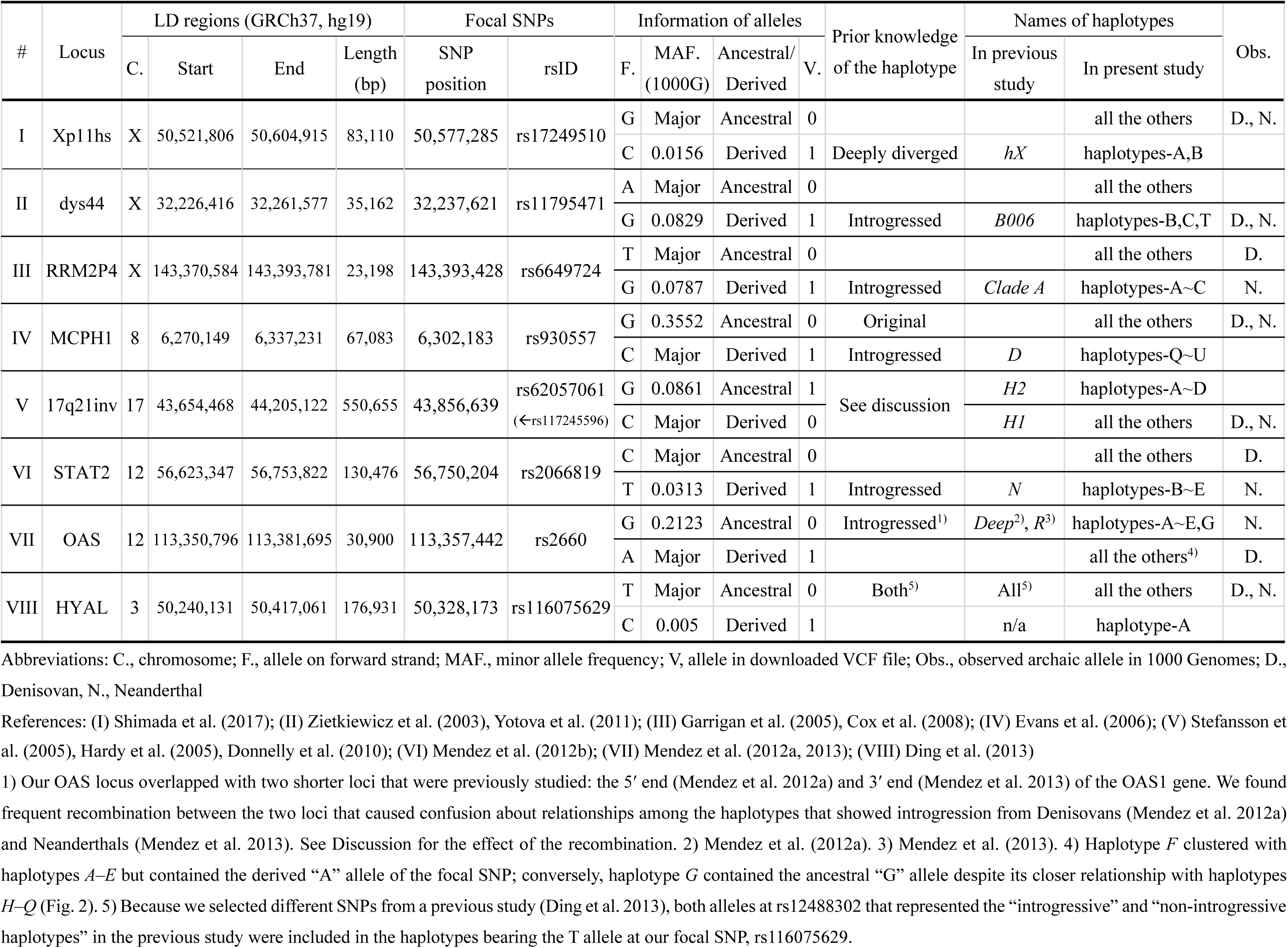
Loci determined by LD region with focal SNPs.

| # | Locus | LD regions (GRCh37, hg19) |  |  |  | Focal SNPs |  | Information of alleles |  |  |  | Prior knowledge of the haplotype | Names of haplotypes |  | Obs. |
| --- | --- | --- | --- | --- | --- | --- | --- | --- | --- | --- | --- | --- | --- | --- | --- |
|  |  | C. | Start | End | Length (bp) | SNP position | rsID | F. | MAF. (1000G) | Ancestral/ Derived | V. |  | In previous study | In present study |  |
| I | Xp11hs | X | 50,521,806 | 50,604,915 | 83,110 | 50,577,285 | rs17249510 | G | Major | Ancestral | 0 |  |  | all the others | D., N. |
|  |  |  |  |  |  |  |  | C | 0.0156 | Derived | 1 | Deeply diverged | *hX* | haplotypes-A,B |  |
| II | dys44 | X | 32,226,416 | 32,261,577 | 35,162 | 32,237,621 | rs11795471 | A | Major | Ancestral | 0 |  |  | all the others |  |
|  |  |  |  |  |  |  |  | G | 0.0829 | Derived | 1 | Introgressed | *B006* | haplotypes-B,C,T | D., N. |
| III | RRM2P4 | X | 143,370,584 | 143,393,781 | 23,198 | 143,393,428 | rs6649724 | T | Major | Ancestral | 0 |  |  | all the others | D. |
|  |  |  |  |  |  |  |  | G | 0.0787 | Derived | 1 | Introgressed | *Clade A* | haplotypes-A~C | N. |
| IV | MCPH1 | 8 | 6,270,149 | 6,337,231 | 67,083 | 6,302,183 | rs930557 | G | 0.3552 | Ancestral | 0 | Original |  | all the others | D., N. |
|  |  |  |  |  |  |  |  | C | Major | Derived | 1 | Introgressed | *D* | haplotypes-Q~U |  |
| V | 17q21inv | 17 | 43,654,468 | 44,205,122 | 550,655 | 43,856,639 | rs62057061<br>(←rs117245596) | G | 0.0861 | Ancestral | 1 | See discussion | *H2* | haplotypes-A~D |  |
|  |  |  |  |  |  |  |  | C | Major | Derived | 0 |  | *H1* | all the others | D., N. |
| VI | STAT2 | 12 | 56,623,347 | 56,753,822 | 130,476 | 56,750,204 | rs2066819 | C | Major | Ancestral | 0 |  |  | all the others | D. |
|  |  |  |  |  |  |  |  | T | 0.0313 | Derived | 1 | Introgressed | *N* | haplotypes-B~E | N. |
| VII | OAS | 12 | 113,350,796 | 113,381,695 | 30,900 | 113,357,442 | rs2660 | G | 0.2123 | Ancestral | 0 | Introgressed<sup>1)</sup> | *Deep*<sup>2)</sup>, *R*<sup>3)</sup> | haplotypes-A~E,G | N. |
|  |  |  |  |  |  |  |  | A | Major | Derived | 1 |  |  | all the others<sup>4)</sup> | D. |
| VIII | HYAL | 3 | 50,240,131 | 50,417,061 | 176,931 | 50,328,173 | rs116075629 | T | Major | Ancestral | 0 | Both<sup>5)</sup> | All<sup>5)</sup> | all the others | D., N. |
|  |  |  |  |  |  |  |  | C | 0.005 | Derived | 1 |  | n/a | haplotype-A |  |

Abbreviations: C., chromosome; F., allele on forward strand; MAF., minor allele frequency; V, allele in downloaded VCF file; Obs., observed archaic allele in 1000 Genomes; D., Denisovan, N., Neanderthal
References: (I) Shimada et al. (2017); (II) Zietkiewicz et al. (2003), Yotova et al. (2011); (III) Garrigan et al. (2005), Cox et al. (2008); (IV) Evans et al. (2006); (V) Stefansson et
al. (2005), Hardy et al. (2005), Donnelly et al. (2010); (VI) Mendez et al. (2012b); (VII) Mendez et al. (2012a, 2013); (VIII) Ding et al. (2013)

## Results

### Gene Genealogies

We constructed a distance-method based phylogenetic tree (i.e., neighbor-joining, NJ) and phylogenetic network for eight loci, and encountered inconsistency in obtained topology between the two methods for four loci: Xp11hs, dys44, MCPH1, and HYAL (Table s1, Fig. 1, Fig. s1, Fig. 2). For example, the allelic genealogy of MCPH1 showed a separated distribution of the haplotypes bearing a derived allele “C” at the focal SNP and specifically discriminated the focal haplotypes, such as haplotype *R* in the network (Fig. 2) despite the clumped distribution in the NJ tree (Fig. 1, see Discussion).

**Figure 1:**
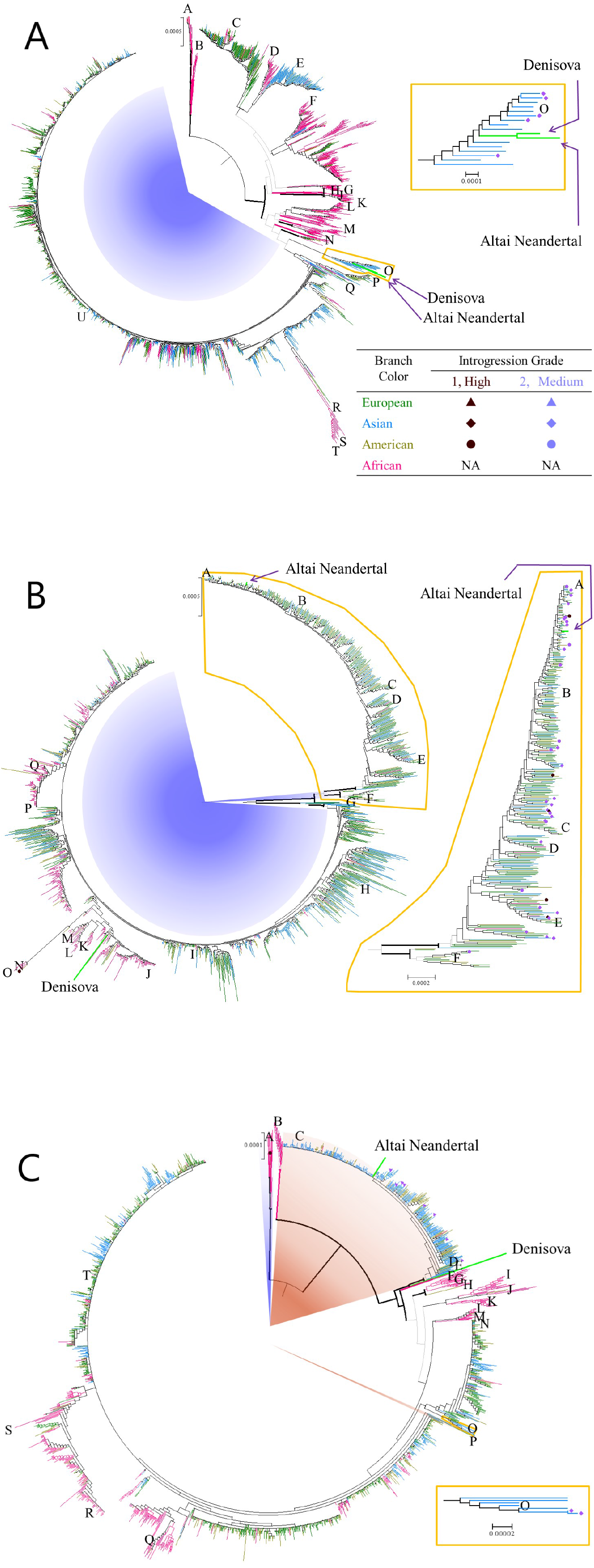
NJ trees for haplotypes of modern humans and archaic hominins (Altai Neanderthals and Denisovans) of three representative loci. (A) MCPH1. (B) OAS. (C) HYAL. Sample origins of haplotypes are expressed by colors of branch tips. Haplotypes of archaic hominins and clusters shared across multiple continents are indicated by light green thick branches and black thin lines, respectively. Line thickness of branches within five bifurcations from the root indicates two classes of bootstrap values of the downward clusters (i.e., less than 50% (thin) and greater than or equal to 50% (thick), respectively). Haplotypes with introgression grades defined by S* analysis are marked by dark red (high) and pale blue (medium). Clusters where one representative haplotype was selected for network analysis are shown in capital letters. Derived allele distributions of focal SNPs in representative haplotypes are depicted by blue background color. When the focal SNP is different from the SNP representing an unusually diverged haplotype reported by the original study, the distribution of the focal SNP from the original study is shown in brown background (see Materials and Methods for details).

**Figure 2:**
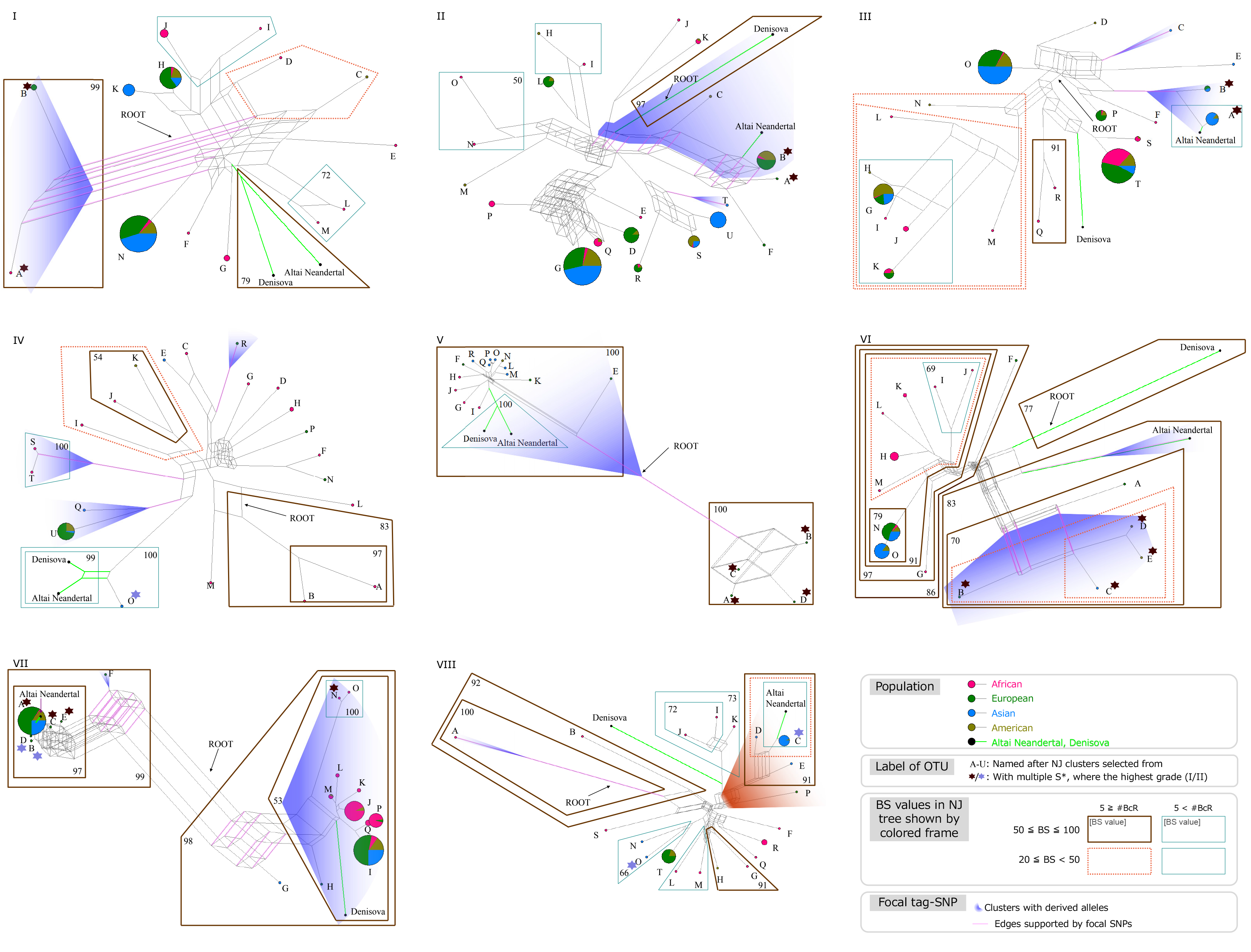
Phylogenetic network of major haplotypes representing major phylogenetic clusters for eight loci. I: Xp11hs, II: dys44, III: RRM2P4, IV: MCPH1, V: 17q21inv, VI: STAT2, VII: OAS, VIII: HYAL. These haplotypes were selected from major clusters of the NJ tree to avoid bias in each locus (see Materials and Methods for details). The color and thickness of frames surrounding haplotypes indicates bootstrap values and distances (i.e., number of bifurcations from the root point) of the clusters in the NJ trees. Distribution of derived alleles of focal SNPs and edges bearing focal SNPs are depicted by the blue area and pink line, respectively.

The phylogenetic networks showed a substantial number of parallelograms in some loci that are characterized by small-sized edges, such as dys44 and HYAL. However, parallelograms with large edges were found in other loci, such as Xp11hs, STAT2, and OAS; this indicates recombination events within a locus (see Discussion).

We classified the haplotype genealogy results into six groups according to tree topological relationships among haplotypes from African, Eurasian, and archaic hominins considering time, place, and direction of gene flow (Fig. 3).

**Figure 3:**
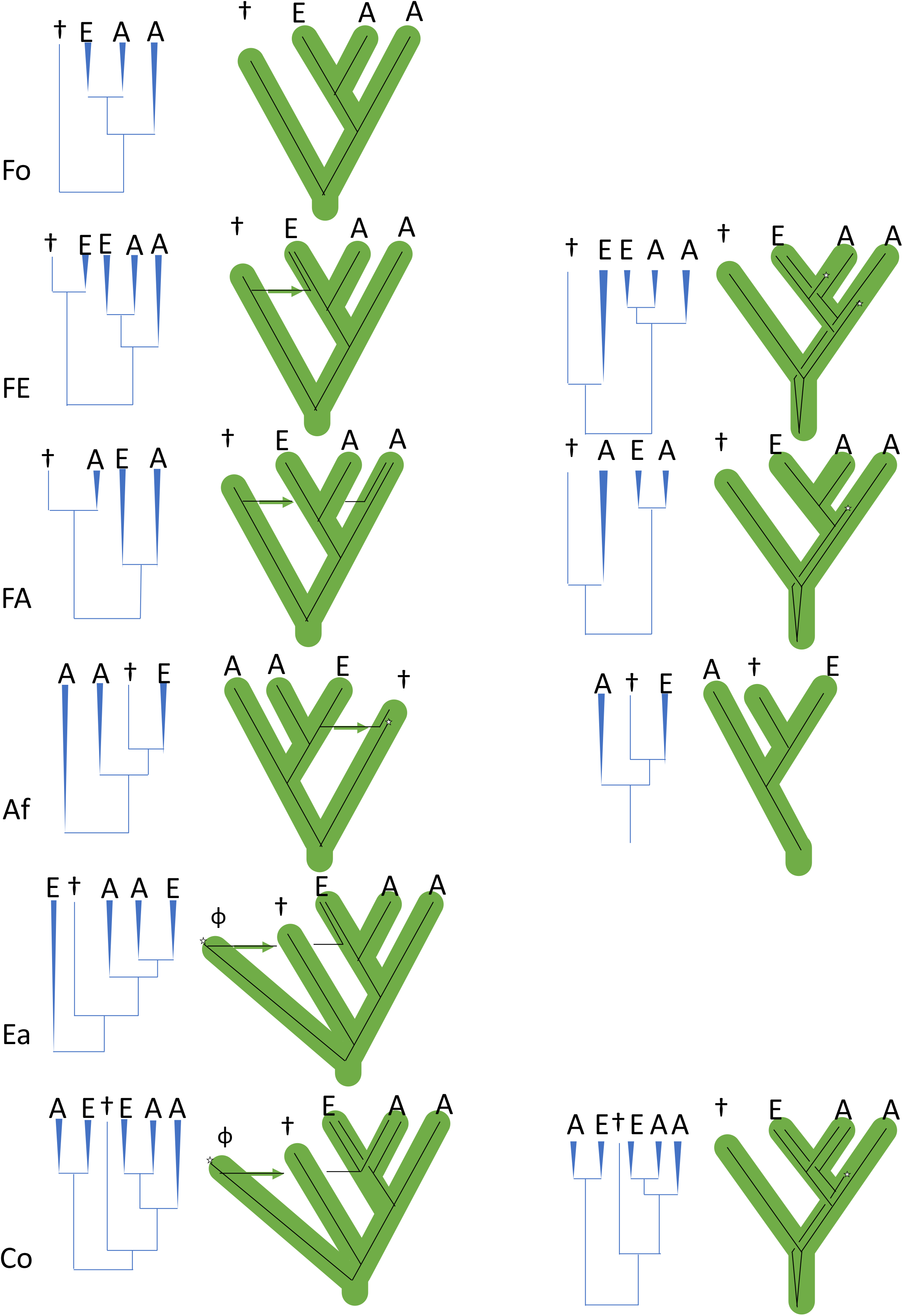
Expected patterns of haplotype genealogy under models without recombination or contamination. A: African cluster, E: Eurasian cluster, †: known archaic haplotype found in Eurasia, φ: unknown archaic haplotype. Type Fo: Recent Out-of-Africa (OOA) without incomplete lineage sorting (ILS); type FE: introgression from known archaic hominins to Eurasians after recent OOA and ILS; type FA: introgression from known archaic hominins to Africa, and ancient polymorphisms within Africa; type Af: introgression from ancestral Eurasians to known archaic hominins, and subdivision within Africa before OOA for both archaic hominins and modern humans (cf., Green et al. 2010, Fig. 6); type Ea: introgression from unknown archaic hominins to Eurasians; type Co: introgression from unknown archaic hominins to an ancestral population prior to OOA, and ancient polymorphism within Africa before OOA for both archaic hominins and modern humans followed by ILS.

For introgression from known archaic hominins (i.e., Altai Neanderthal and Denisovan in this study) to modern humans, we considered the possibilities of post-OOA in Eurasia [type FE] and pre/post-OOA in Africa [type FA]. We did not distinguish pre- and post-OOA introgressions that have remained within Africa, because they were expected to be indistinguishable in haplotype genealogy. Alternatively, introgression from ancestors of modern Eurasians to archaic humans was expected to have occurred after OOA in Eurasia [type Af]. We also considered the possibility of introgression from unknown archaic hominins to modern humans, which occurred both post-OOA in Eurasia [type Ea] and pre-OOA in Africa [type Co].

We could not rule out the possibility of ILS of ancestral polymorphisms by haplotype tree topology for three types [types FE, FA, and Co] within these classifications.

### S*

S* is a method that enables estimation of the presence and amount of gene flow between sub-populations by detecting combinations of rare alleles (Plagnol and Wall 2006; Vernot et al. 2016). We performed S* analysis to estimate distinct gene flow events from archaic hominins after OOA using Africans as a reference population and detect novel SNP allele combinations in modern humans that only exist in Eurasia. We modified S* analysis to apply phased massive genome sequence data and highlight haplotypes with high S* scores. Then, we classified obtained S* into three classes (i.e., high, medium, and low introgression grades; see ‘S* analysis’; ‘Algorithm’ in Materials and Methods). The S* score showed signs of gene flow after OOA in all eight examined loci to a greater or lesser degree (Table 3, Fig. s2). We observed clusters that included haplotypes with high S* scores and haplotypes of known archaic hominins in clusters *A* and *B* at dys44, clusters *A* and *B* at RRM2P4, cluster *O* at MCPH1, clusters *A* to *F* at OAS, and clusters *C* and *O* at HYAL (Table 3, Fig. 1). These findings indicate the possibility of gene flow between the archaic hominins and Eurasians after OOA. Three of the loci (dys44, RRM2P4, and OAS) showed type FE topology, which supports introgression in Eurasia after OOA (Fig. 1, Table 2).

**Table 2.**
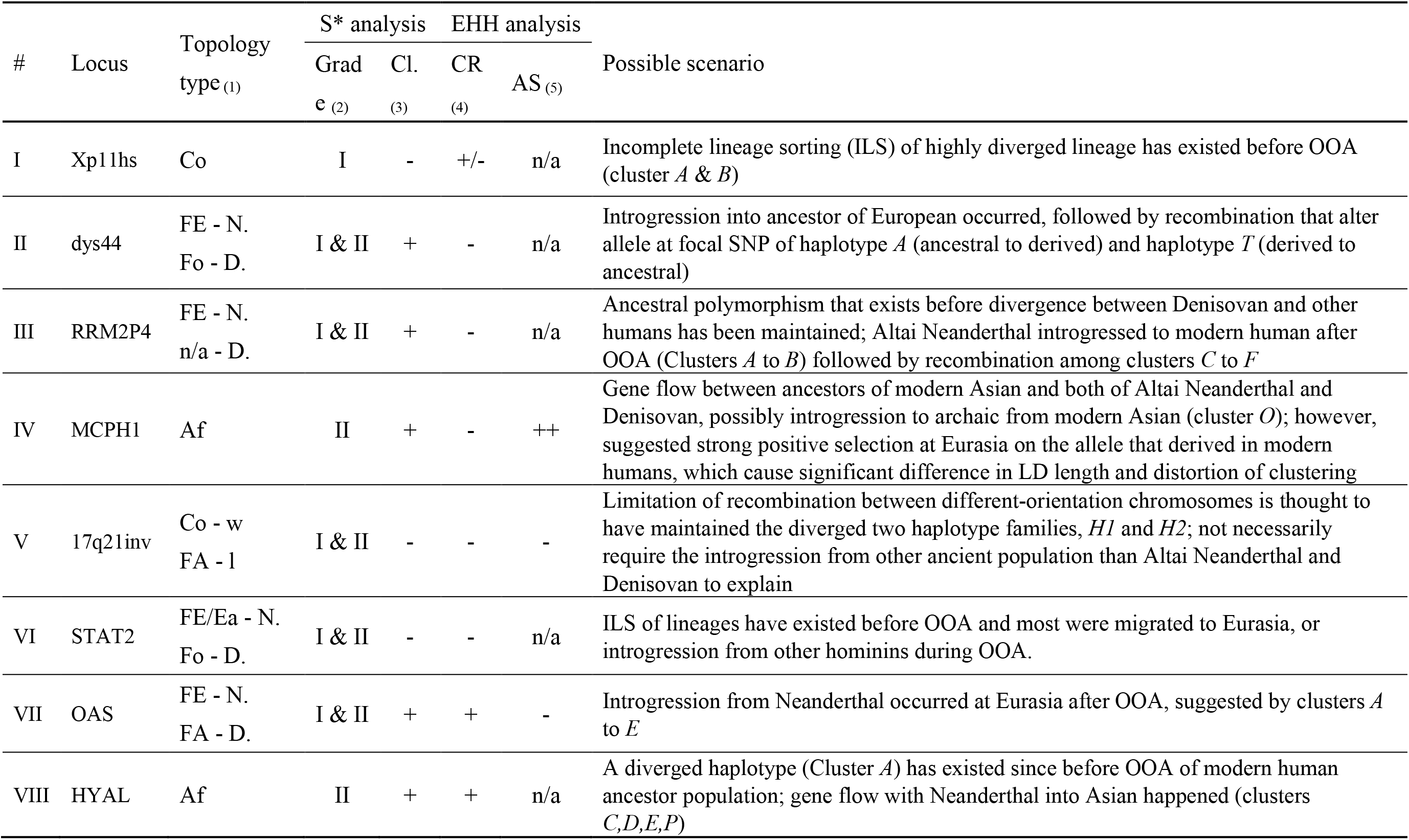
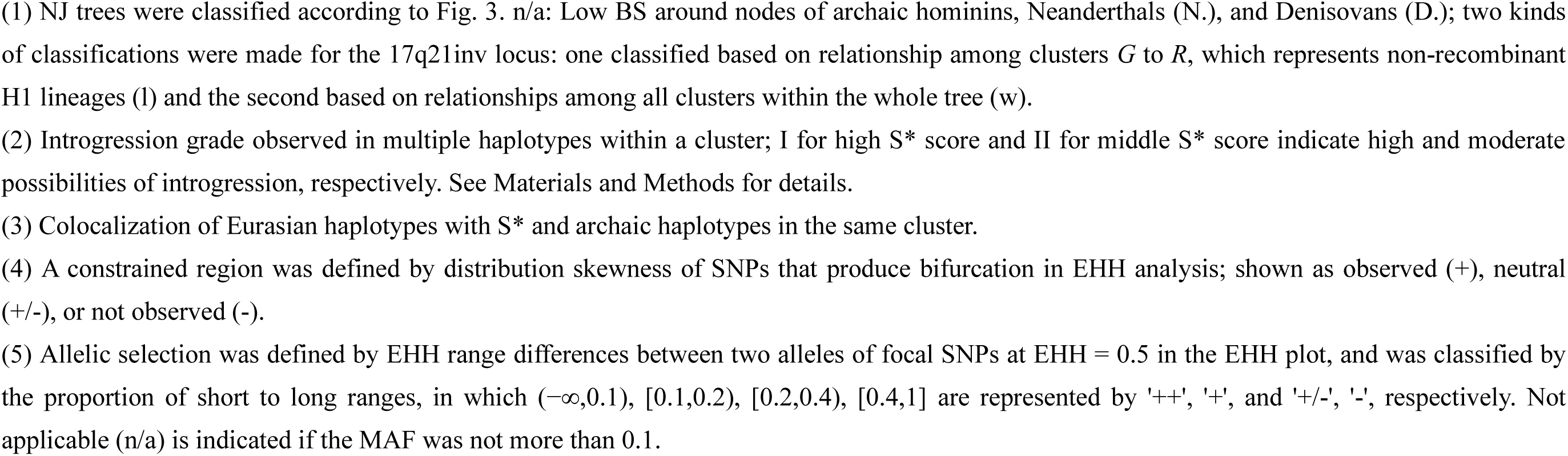
Summary of results

**Table 3.** S* analysis results

| # | Locus | Tree topological relationship of modern human |  | Clustering pattern of haplotypes with S* | Miscellaneous note |
| --- | --- | --- | --- | --- | --- |
|  |  | to Neanderthal | to Denisovan |  |  |
| I | Xp11hs | Closely related with an African cluster | Closely related with an African cluster | Except five African haplotypes used as reference, all haplotypes in the outmost cosmopolitan cluster marked high-grade S* | No S* was observed other than the outmost cluster (*A* and *B*) |
| II | dys44 | Clustered with European | External and Independent | Cosmopolitan outer cluster includes high-grade S* and Neanderthal | High S* independently marked on haplotype *T* that focal SNP allele suggest to be a recombinant |
| III | RRM2P4 | Clustered with Eurasian | Independent | Cosmopolitan outer cluster includes high-grade S* and Neanderthal | Clusters *C* to *F* suggests another but related event with introgression from Altai Neanderthal |
| IV | MCPH1 | Clustered with S* Asian | Clustered within Asian with S* | Inner cluster includes multiple Asian with medium-grade S*, Neanderthal, and Denisovan | Strongly suggested gene flow between Asian and archaic humans |
| V | 17q21inv | Clustered with African | Clustered with African | Outmost cluster includes *H2* haplotypes with high or medium-grade S* | Reference population may not contain enough number of Africans with inverted haplotype *H2*, which resulted in high S* scores in *H2* family |
| VI | STAT2 | Independent | Independent | Outer cluster includes multiple Eurasian with S* and located closer to Neanderthal and Denisovan than other modern human clusters | S* suggests introgression from unknown but related with Neanderthal or Denisovan |
| VII | OAS | Clustered with S* Eurasian | Closely related with inner African clusters | Cosmopolitan outer cluster including high-grade S* and Neanderthal | S* on haplotype on an American clustering with African may suggest novel combination of rare alleles by recent recombination (haplotype *N*) |
| VIII | HYAL | Clustered with Eurasian | Independent | Eurasian cluster *C* includes Neanderthal and multiple medium-grade S*; outmost African cluster *A* includes one American with high-grade S* | S* at cluster *O* can be recombination according to network topology |

Clusters composed of archaic hominins and Eurasians also had high S* scores in cluster *O* in MCPH1 and cluster *C* in HYAL loci, but these topologies were not type FE but type Af (Table 2, Table 3). The S* scores in these clusters suggested one of the two Af scenarios: ancient subdivision within Africa before the leaving of Neanderthals from Africa (Fig. 3, right of Af, See Discussion). In the HYAL locus, cluster *O* showed a medium S* score in a small number of haplotypes. Moreover, the position of cluster *O* in the network suggested recombination between *N* and *T*, which may produce SNP combinations not found in the reference population (Africans).

We also observed S* haplotypes in the outermost cluster that were located close to but separate from the haplotypes of known archaic hominins in the gene genealogies of Xp11hs and STAT2 (Table 3). These diverged clusters of Xp11hs and STAT2 contained high S* haplotypes composed of various populations (cosmopolitan clusters) without geographically aggregated sub-clusters. Generally, a random geographic distribution is considered ILS (Zhou et al. 2017), and these diverged clusters in these two loci can be attributed to events that produce polymorphisms that existed before or during OOA, rather than introgression from archaic humans after OOA. Although a similar pattern was also shown in the 17q21inv locus, introgression was not necessarily needed to explain this pattern because of limited recombination between chromosomes, with different orientations caused by inversions (see Discussion).

Some of these phylogenetic trees and networks showed the effect of recent admixture in Americans, because American samples used in the 1000 Genomes Project are “admixture individuals” from various North Americans, not native American individuals, which is documented as “Ad Mixed American” in The International Genome Sample Resource (http://www.internationalgenome.org/faq/which-populations-are-part-your-study) (Table s2). Two American haplotypes with high S* scores were observed in African clusters (cluster *A* at HYAL and cluster *N* at OAS). Both of these African clusters were small and separated from other African clusters in the tree. These findings indicate that Americans inherited these haplotypes from Africans, and these haplotypes are even rare in Africans, which resulted in high and medium S* scores on these haplotypes; therefore, these S* scores may not necessarily be caused by introgression.

### Extended Haplotype Homozygosity

Extended Haplotype Homozygosity (EHH) is a measurement that indicates the probability of the presence of a continuous linkage disequilibrium (LD) block and is defined as the probability that two randomly chosen chromosomes bearing the same allele of a given focal SNP site are identical haplotypes within a genomic region that is *x* distance from the focal SNP site (Sabeti, PC et al. 2002). EHH was developed to detect positively selected alleles through comparison of LD block presence probability of focal SNP alleles. We used EHH to evaluate our locus-defining method that was determined by LD *r^2^* measure.

Comparison of genomic region length ratio of EHH to LD (*R.length*) indicated that EHH regions were shorter than LD regions in all examined loci (Fig. 4a, Table s3). Although the *R.length* ranged two orders of magnitude (0.005–0.462), the SNP density ratio of EHH to LD regions showed a 1.57-fold difference (0.856–1.345; Fig. 4b, Table s3); this indicated that a smaller EHH than LD is not caused by the poor availability of SNP data. We suggest that the *R.length* difference resulted from differences in the extent of recombination (Table s3).

**Figure 4:**
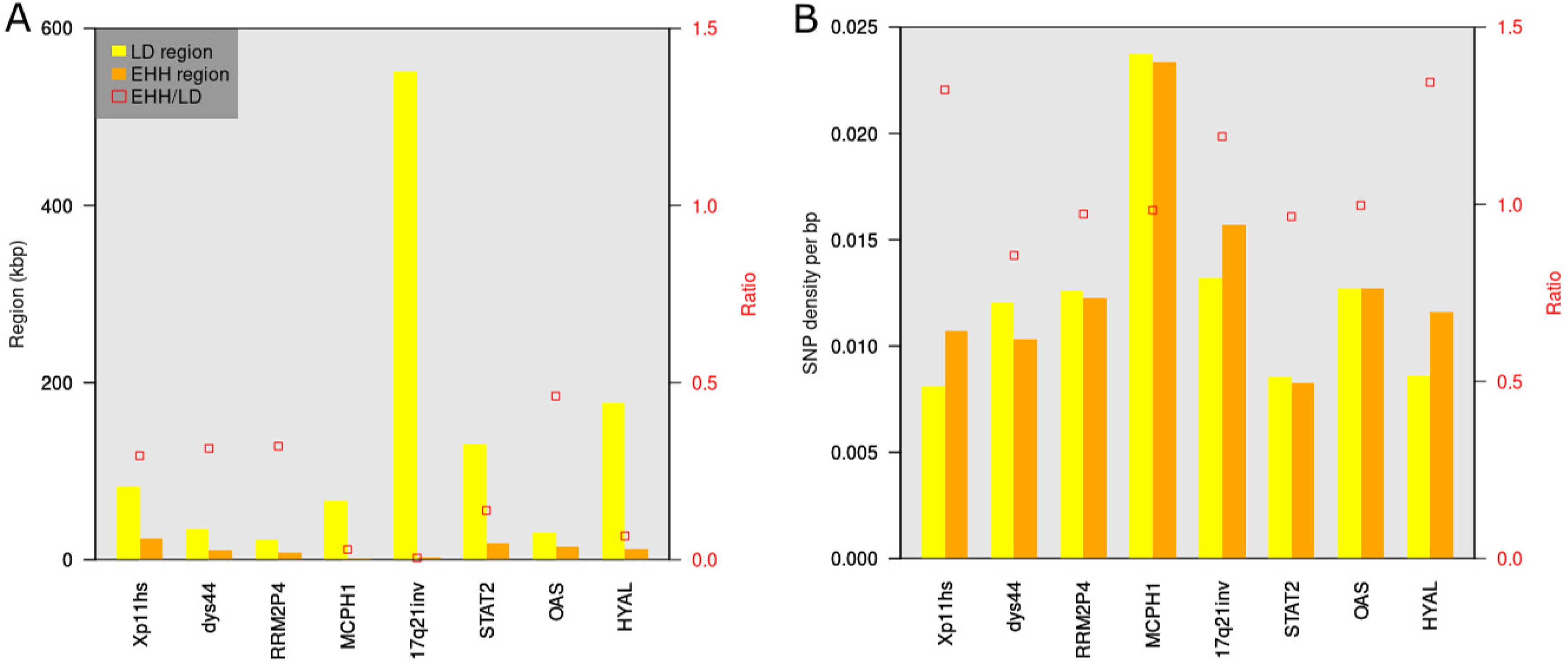
Comparison between LD and EHH regions. Comparison of (A) length and (B) SNP density. Bars for regions and red plots for EHH/LD ratio graphed against left and right vertical axis, respectively.

The bifurcation graphs of EHH analysis showed bifurcations of multiple lineages at a single SNP position, which suggests exchange of SNP alleles between haplotypes via recombination (Fig. 5).

**Figure 5:**
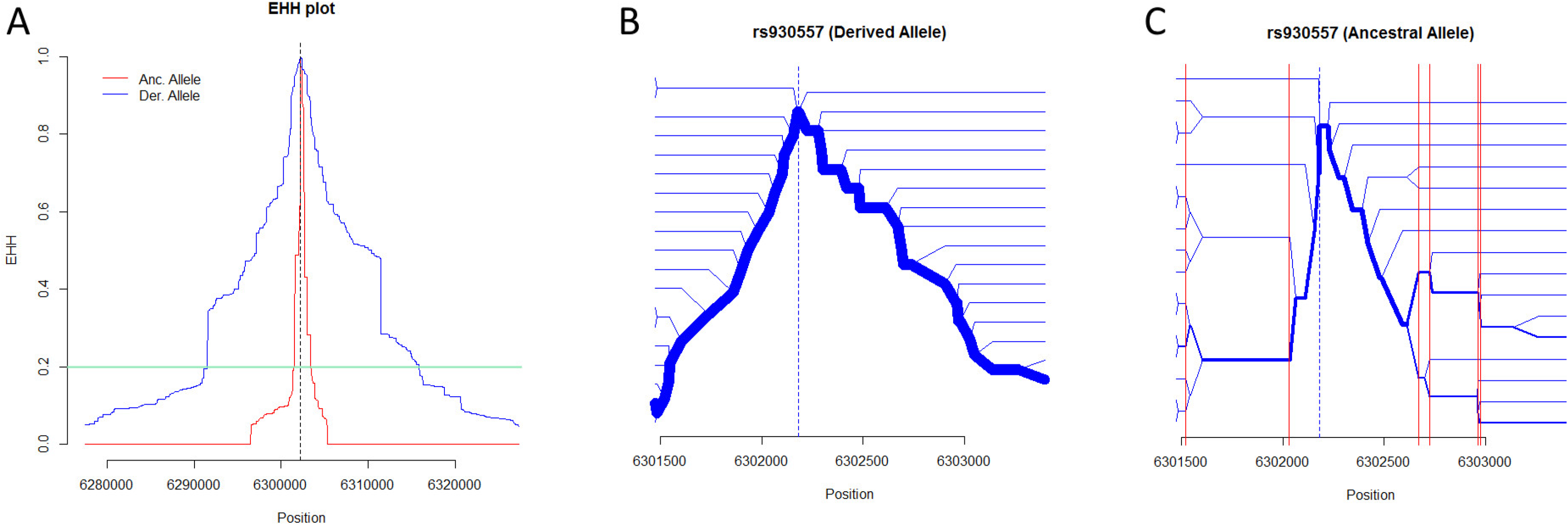
EHH analysis for MCPH1. (A) EHH plot of MCPH1. EHH are plotted in the genomic region showing EHH < 0.05 in at least one allele. Red and blue lines indicate EHH for ancestral (Anc) and derived (Der) alleles, respectively. The bifurcation graphs were generated within the region showing EHH > 0.2 (aqua green line) in both alleles. (B, C) Bifurcation graphs for the MCPH1 locus. The position of the focal SNP site is shown by blue dotted lines. The width of blue lines represents the frequency of haplotypes bearing derived (B) and ancestral (C) alleles of each focal SNP. Red lines in (C) indicate SNP positions that make bifurcations at multiple branches. EHH analysis of other loci is shown in Supplementary Figure s4.

Our EHH analysis indicated selection on only a specific allele in the MCPH1 locus among the eight examined loci (Table 2 EHH column, Table s3). The EHH range of derived allele “C” from rs930557 was longer than that of ancestral one “G,” which is explained by selective sweep in the MCPH1 locus (Fig. 5a). The MCPH1 bifurcation graph for ancestral allele “G” showed succession of bifurcations in multiple branches at common genomic positions, such as 6301472, 6301546, 6302671, 6302962, and 6302971, which indicates the existence of SNP sites that share alleles with other haplotypes (Fig. 5b & 5c, Fig. s3). This indicates the existence of recombination among haplotypes bearing the ancestral allele “G” at rs930557 (Fig. 5c). Meanwhile, however, almost all bifurcations were observed in a single lineage of haplotypes with the derived allele “C” at rs930557, which suggests that a novel mutation generated a novel bifurcation (Fig. 5b). This is explained by selective sweep of haplotypes bearing the derived allele “C”.

Although Stefansson (2005) suggested positive selection of the H2 lineage in the 17q21inv locus, this was not confirmed by our EHH (Table 1).

We also noted that the bifurcation graphs depicted a skewed distribution of branching points of haplotypes in OAS, HYAL, and Xp11hs (Fig. s4). Consequently, the numbers of haplotypes did not increase in proportion to the distance from the focal SNP position in these regions. This is due to SNP density change, which represents the existence of differences in evolutionary constraints within the EHH region; the region with fewer SNP sites was confirmed to overlap with the promoter region of the SHROOM4 gene in Xp11hs, transcribed region of the OAS1 gene, and the transcribed region of the HYAL3 gene (Table 2, column “CR,” Table s3). Thus, EHH analysis indicated the existence and extent of selective pressure on haplotypes.

## Discussion

### Significance of This Study

This study demonstrated that each locus has their own evolutionary history, which was previously missed by allele frequency-based analyses. Using massive individual genome data, our combinatorial analysis that consisted of tree topology classification, S*, and EHH analyses can clarify how selection pressure varies by haplotype. We demonstrated the effectiveness, utility, and reliability of each analysis. First, in haplotype genealogy, clustering of archaic hominins with multiple modern humans with high S* scores likely represents introgression. Second, EHH analysis is useful for detecting regions that are under functional constraint and selective sweep. The length ratio between LD and EHH regions is useful for clarifying the amount of recombination. We also identified concepts that should be discussed further, such as comprehensiveness of African samples and definition of loci in genome-wide individual genome sequence data that may contain various recombinations in different genomic locations depending on haplotypes.

### Signs of Introgression from Archaic *Homo* and Diversity in *H. sapiens*

This study confirmed multiple hybridization events caused by divergence followed by subsequent contact after isolation. Introgression from archaic hominins is highly likely when a haplotype of archaic hominins is clustered with multiple modern haplotypes with high S* scores. However, high S* scoring modern haplotypes without clustering with archaic hominins may have been caused by insufficient samples sizes of the reference population, novel combinations of rare alleles by recent recombination, and gene flow with unknown archaic hominins.

An earlier study showed several gene flow events among archaic human groups, which included unknown archaic groups (i.e., not Neanderthals and Denisovans) (Prufer et al. 2014). This study demonstrated that *H. sapiens* experienced more population subdivision and hybridization events than expected based on known introgression from Neanderthals and Denisovans. ILS of ancestral polymorphisms alone cannot explain the complexity that we showed. Our findings indicate the presence of highly structured populations within Africa, which includes ILS during population subdivision and several introgression events with unknown populations of genus *Homo*.

The Neanderthal and Denisovan haplotypes had different locations in the tree topology of three loci (dys44, RRM2P4, OAS), and the Neanderthal haplotypes showed the possibility of introgression (type FE), but those of Denisovans did not show a clear trend (Table 2). This difference between Neanderthal and Denisovan haplotypes resulted from a history of migration and hybridization with modern humans. It is noteworthy that Denisovan genomes contained components that were introgressed from other archaic populations, which were deeply diverged from a common ancestor of Neanderthal, Denisovan, and modern humans (Prufer et al. 2014).

### Effects of Selection

As the 1000 Genomes Consortium observed, rare variants generally originate by recent mutation, which causes a negative correlation between variant frequency and haplotype length (The 1000 Genomes Project Consortium 2012). As expected from this relationship, comparatively longer EHH were observed in rare alleles with minor allele frequency (MAF) < 0.1 in bifurcation graphs of our EHH analyses (Table s3), we did not further analyze these rare alleles. Without considering this relationship, an apparent longer EHH of a rare allele compared with that of a major allele may produce a misleading inference about the selection of haplotypes with rare alleles. The phylogeographic network of the HYAL locus indicates that the most diverged haplotype A accumulated a lot of singleton SNP variants, although only small parts of SNP sites were shared with haplotypes B and S, which indicates limited recombination among them (Fig. 2). The EHH analysis did not provide enough evidence for selection for the haplotype A because of a low frequency of the minor allele carried by haplotype A (Table s3, Fig. s4). Considering branch lengths and phylogenetic relationships including ancient genomes of other hominins, haplotype A of the HYAL locus may be a rare neutral variant that existed in modern humans in Africa before the divergence of the modern human lineage from archaic human groups such as Neanderthals and Denisovans.

### Effects of Recombination

Some of the median networks in this study formed complex aggregation of parallelograms (Table s1, column ‘Size and frequency of reticulation in phylogenetic network’). We manually omitted a considerable amount of parallelograms (see Materials and Methods) because of the large sample size and long locus regions. Because we constructed median networks that were categorized as split networks, consecutive SNPs in genome position observed on parallel edges implicitly represent evolutionary events that occurred on a genomic fragment, such as recombination, horizontal gene transfer, or gene duplication (systematic error) (Huson and Bryant 2006); this is more likely for large parallelograms that have long edges with numerous consecutive SNPs. Alternatively, short edges with a small number of SNPs separated from each other can be formed by parallel and convergent substitution. Huson and Bryant (2006) distinguished the systematic error from the sampling error, which is random error that results from a small sample size (number of SNP sites). Then, they highlighted that the rapid growth in availability of large genomic sequences increased the importance of systematic errors but diminished the impact of sampling error on phylogenetic inference. Accordingly, we actually found that recombination resulted in systematic errors such as large parallelograms in our phylogenetic networks, especially because we defined a locus being as long as possible by LD in this study. However, owing to our definition of a locus, we observed recombination between the two short loci used in separate two studies that determined haplotypes of the OAS gene region (Mendez et al. 2012a; 2013). Mendez *et al*. (2012a) determined haplotypes based on the 5’ end region, which included exons 1–3, whereas Mendez *et al*. (2013) started typing based on 15 SNPs that spanned about 760 bp at the 3’ end, which included exons 4–6 of the OAS1 gene. The recombination between the two short loci may cause confusion about relationships among the haplotypes determined by the two studies, because genealogical relationships among haplotypes are recognized by landmark haplotypes, such as haplotypes of the human reference genome, the two archaic humans, and introgression candidates. Our locus (30.9 kb) determined by LD block overlapped with the 3’ end; this included exons 4–6 of the OAS1 gene, which were the focus of the study conducted by Mendez *et al*. (2013). Consequently, our results were consistent with those of Mendez *et al*. (2013) and showed that Eurasian haplotypes had a close relationship with Neanderthals (topology type FE), although Mendez *et al*. (2013) used Neanderthals from Vindija Cave, Croatia, whereas we used a Neanderthal from the Altai Mountains, Russia. Even though Mendez *et al*. (2013) did not explicitly discuss the relationship with Denisovan haplotypes, our results based on the overlapping genomic region with Mendez *et al*. (2013) represent a distant relationship between Denisovans and Neanderthals (Fig. 1, Fig. 2). Our study did not provide evidence of post-OOA introgression from Denisovans (topology type FA, Table 2), although our loci are included in their “Denisova Introgressive Block (~90 kb)” that was introgressed from Denisovans to Melanesians, which was detected using the HGDP panel mentioned in the Mendez *et al*. (2012a) and depicted in Figure 1 of Mendez *et al*. (2013). This difference in results is probably because Melanesians were not included in our samples. Further investigation of recombination between the two loci that includes Melanesian samples is needed.

Recombination also potentially affects S* analysis. Because S* score indicates the possibility of introgression based on two rare alleles that are colocalized within a haplotype that are absent from the reference population (Africans in this case), recombination may generate novel combinations of two rare alleles in a recombinant haplotype that yields a high S* score. In this study, we found one candidate of this example in haplotype O of the HYAL locus (Fig. 1).

### Inconsistency between Phylogenetic Inferences by Network and Distance Methods

Our close examination provides a rationale for the differences in inferences of phylogenetic relationships between the network based on character data and the tree that was constructed using distance methods (i.e., the NJ method). This was observed for the MCPH1 locus. A single mutational event can explain the allele distribution of the focal SNP rs930557 on the NJ tree but not the phylogenetic network (Fig. 1, Fig. 2). That is, haplotype *R* is separate from haplotypes *Q, S, T*, and *U*. The rationale of this contradiction can be explained as follows. First, the difference in selection pressure among haplotypes likely produced differences in LD length among them (Table s3). Then, the LD lengths of haplotypes with beneficial alleles became longer because of less frequent recombination than other haplotypes; EHH of the MCPH1 locus indicated that this is a selective sweep (Fig. 5a). Because the recombinant haplotypes carry a mixture of different ancestral information, the numbers of SNP sites that shared ancestry (synapomorphic SNPs) in the examined loci were inconsistent among haplotypes; that is, haplotypes with short EHH shared fewer synapomorphic SNPs than those with long EHH. This might produce systematic error when inferring phylogeny among haplotypes of a locus. Therefore, this inconsistency in evolutionary background among haplotypes results from the definition of loci that were uniformly determined by their *r^2^* values. Second, a phylogenetic network based on character data is thought to be more vulnerable than the distance method to inconsistency in LD length, because distance methods include a correction process with substitution models study (Kishino and Hasegawa 1989; Felsenstein and Churchill 1996), such as the F84 model in the present; this differs from the median network, which is based on character state without any weighting for synapomorphic SNPs.

### Polymorphic Inversion

Among the loci in this study, the genomic region with a 900-kb inversion polymorphism at 17q21.31 (17q21inv) had the most abundant accumulation of knowledge from previous studies. Previous studies showed that the inversions are found in a region where recombination was not observed around 2 Mb (Evans et al. 2004; Oliveira et al. 2004; Pittman et al. 2004; Fung et al. 2005). The 17q21.31 region has been characterized as a region that is rich in chromosome rearrangements that are accompanied by segmental duplications (SDs), which frequently and repeatedly occurred during primate evolution (Zody et al. 2008). SDs play a critical role in chromosomal rearrangement during primate evolution (Bailey and Eichler 2006) (e.g., Shimada et al. 2005).

Because of frequent evolutionary changes, defining ancestral haplotype is not simple, and the evolutionary history of 17q21inv is still under debate (Alves et al. 2012; Steinberg et al. 2012). Zody et al. (2008) showed that the 17q21inv polymorphism is specific to the human lineage. Baker et al. (1999) named the common (non-inverted) haplotype H1 and the rare (inverted) haplotype H2. According to the model proposed by Steinberg et al. (2012), the inverted orientation (H2 haplotype) was the ancestral state of the *Homo* lineage, and was replaced by the H1 haplotype, which emerged by (re-)inversion approximately 2.3 million years ago (Mya). This predominance of the H1 haplotype is supported by the observation of this haplotype in Neanderthal (Green et al. 2010) and Denisovan genomes (Setó-Salvia et al. 2012), which was also supported by our results. Two studies that evaluated population genomics using SNP genotype data from worldwide populations focused on H1 (Steinberg et al. 2012) and H2 (Alves et al. 2015) haplotype families, and found that these haplotypes independently had African clusters that diverged first within each haplotype family. This indicates that both H1 and H2 haplotype families existed within Africa before OOA of modern humans (*H. sapiens*). Lack of reinforcement of African samples in our study may explain why the H2 haplotype family (haplotypes *A–D*) did not form a cluster that only consisted of Africans in our phylogenetic analysis. Therefore, these previous studies supported the idea that ancestral polymorphisms were maintained before the divergence of modern humans from the ancestral Neanderthals and Denisovans (topology types FA and Co, Fig. 3). Those previous studies also showed that copy number polymorphism of the SDs arose in the H1 and H2 lineages around 250,000 years ago and 1.3 Mya, respectively, and named the haplotypes based on haplotype family and presence/absence of SDs. For example, H1’ and H1D represent haplotypes without and with SDs of the H1 lineage, respectively. Alves et al. (2015) showed that North Africans have more H2D than H2’ that is closer proportion to non-Africans, but rare H2D in Sub-Saharan Africa in which H2’ form deepest monophyletic clade; this indicated that H2’ was maintained within Sub-Saharan Africa during OOA of modern humans. Furthermore, Alves et al. (2015) also demonstrated negative trends of Tajima’s D (Tajima 1989) in both H1 and H2 lineages, although H1 is more variable than H2 in nucleotide diversity. This deviation from neutrality for both the H1 and H2 lineages challenge the possibility of selective sweep on H2, although the authors carefully discussed that this was a speculative suggestion, and they did not specify if demography or positive selection was the cause of current geographic patterns.

Generally, a demographic event does not affect only a single locus. Our EHH analysis does not suggest a selective sweep in the H2 lineage or any notable difference between the two haplotypes. Considering restrictions in recombination between inverted and non-inverted haplotype families, the negative trends of Tajima’s D can be explained by each haplotype family acting like a genetic barrier, which divided haplotypes and resulted in a smaller effective population size and longer LD than other loci; this likely indicates a population just after admixture of two divided populations. Our study indicated that the current distribution of H2 haplotypes is irrelevant to contact with Neanderthals and/or Denisovans. Additionally, we suggest that introgression from other unknown ancestral humans is not necessarily required to explain haplotype distributions at the 17q21inv locus. Consequently, as a more likely scenario, long-lasting ancestral polymorphisms with restricted recombination between the two haplotype families probably resulted in the topology type Co haplotype phylogeny, although one basal bifurcated cluster (H2 cluster) only consisted of Eurasia, probably because of the loss of haplotype variation from Africa and archaic humans or insufficient sampling.

### Distinguishing Introgression and ILS

To date, efforts have been made to distinguish introgression after hybridization and ILS of ancestral polymorphisms (Joly et al. 2009; Kubatko 2009; Meng and Kubatko 2009; Green et al. 2010; Gerard et al. 2011; Nakhleh 2013; Yu et al. 2014; Martin et al. 2015; Zhou et al. 2017; Edelman et al. 2019; Kubatko and Chifman 2019). Those studies evaluated tree topology, divergence time, and geographic distribution of alleles/haplotypes (Table s5). Based on those lines of evidence, these prior studies represented three approaches.

The first is a genealogy-based approach. This approach was used by Joly et al. (2009), who focused on the relationship between gene trees and species/population tree. In particular, the expectation of minimum divergence between two haplotypes is smaller for the hybridization model than that for ILS (Joly et al. 2009, Fig1). Through simulation and empirical application, they demonstrated that the detection power of hybridization is reduced in larger population sizes and shorter sequences; this provided incentive for our comparison among loci that were as long as possible based on the common data set of the 1000 Genomes Project (see ‘How to Treat Locus’).

The second approach is an allele frequency spectrum-based approach that focused on relative allele frequency of shared derived alleles in four taxa, and uses D statistics or ABBA test (Green et al. 2010; Martin et al. 2015).

The third approach is a geographic information-based approach that uses information about habitat changes during subdivision and migration of populations/species to estimate gene flow. Estimates of historical change of habitats using ecological modeling tools (e.g., MaxEnt; Elith et al. 2011) are combined with estimates of demographic modeling performed by the coalescent-based isolation-with-migration model (Hey and Nielsen 2004) or admixture analysis using STRUCTURE (Hubisz et al. 2009), which is typically used to compare sympatric and allopatric populations (e.g., Zhou et al. 2017).

The present study proposes that haplotype-based S* analysis combined with categorization of tree topology is a simple and model-free method to identify introgression from ancient humans with fewer assumptions. It is assumed that Eurasians must be a subpopulation of Africans (a reference population) under the OOA model; this means that all original or closely related haplotypes of the OOA population (Eurasian) should remain in Africa today and included in the reference population of S* analysis through vast sampling with a sufficient sample size. If these conditions are not fulfilled, a false positive S* signal might occur.

Haplotype tree topology alone cannot distinguish ILS and introgression, as observed in tree types FE, FA, and Co (Fig. 3). However, the Eurasian haplotypes with high S* scores clustered with ancient haplotypes, which strongly suggested introgression from ancient humans in the case of type FE (Fig. 3); this may be applicable to Neanderthal branching in loci dys44, RRM2P4, and OAS (Fig. 1). Although the Neanderthal haplotype first coalesced with the high S*-score cosmopolitan cluster in STAT2, the Neanderthal haplotype was not included in the cluster, as shown in other loci classified as type FE (Fig. 1). This could be explained by introgression occurring just after OOA before divergence between Europeans and Asians if the assumption regarding sample size of reference population is fulfilled. If the assumption is not fulfilled, a high S* score in the reference population that underwent ancient subdivision in Africa can produce a false positive regarding absence of African sister haplotypes of the Eurasian haplotypes. Because Eurasians are not involved with introgression focused in discrimination within type FA, our application of S* analysis does not distinguish between “introgression from known archaic to modern Africans” (Fig. 3, left of FA) and “ILS of ancient polymorphisms within Africa” (Fig. 3, right of FA).

In the outermost clusters A and B of the locus Xp11hs, Eurasian haplotypes with high S* scores clustered together with the five haplotypes belonging the African reference population (type Co, Fig. 3). This aberrant clustering of S* and reference haplotypes might be explained by these five African haplotypes representing a small proportion of the all 377 African haplotypes used as reference in S* calculation. The individuals of the five African haplotypes included both East and West Africans (Tables s6–8); this may indicate that ancient polymorphisms persisted before subdivision between East and West Africans. Further simulation-based study that focuses on these two scenarios is needed for this tree type.

A high S* score at the basally diverged Eurasian lineage under the topology type Ea clearly indicates introgression from unknown archaic hominins in Eurasia (Fig. 3). This pattern was partially found in the locus STAT2 and is consistent with the following published scenario that explains an observation that African genomes shared more derived alleles with the Neanderthal genome than with the Denisovan genome: “Denisovans received gene flow from ancestors that were deeply diverged from common ancestors among Neanderthals, Denisovans, and modern humans” (Prufer et al. 2014).

The gene tree type Af under the conventional population tree [(archaic, (South African, (East African, Eurasian)))] indicates introgression from ancestral Eurasians to known archaic hominins (Fig. 3, left of Af). If the conventional population tree is not assumed, another scenario can be considered for type Af: ancient subdivision within Africa before the leaving of Neanderthals from Africa (Fig. 3, right of Af). This ancient subdivision model assumes ancient population structure that persisted before divergence of the ancestral Neanderthal population until OOA of modern humans (Fig. 3, right of Af; cf., Green et al. (2010), Fig. 6; Wall et al, (2013), Fig. 1a). This model is not supported by previous studies (Sankararaman et al. 2012; Yang et al. 2012; Wall et al. 2013). In this study, the MCPH1 and HYAL loci were classified into type Af. Although low stability of relationships among major clusters prevents a conclusive statement (Table s1), cluster *O* in the MCPH1 gene tree revealed that Neanderthal and Denisovan haplotypes exhibited monophyly and exclusively clustered with modern Asian (Fig. 1). Because the Neanderthal sequence originated from the Altai Mountains, which are geographically close to the sampling location of Denisovans, cluster *O* indicated gene flow between ancestral modern Asians and these two ancient hominins within a limited area of Asia. The direction of gene flow depends on the assumed population tree or model. Given the conventional population tree model, gene flow from modern Asians to these archaic hominins is reasonable. However, reverse gene flow was concluded under the ancient subdivision in the Africa model. This consideration necessitates further empirical studies.

As discussed above, S* analysis based on gene trees is valuable for distinguishing ILS and introgression in at least some cases. However, researchers must be cautious when selecting reference populations. Our modification of S* based on rare/minor alleles assume that the African population represents universal human variation. Thus, insufficient sampling from African populations and extinction of ancestral African populations produces false positive S* scoring, which may especially occur when highly diverged population structure existed before OOA. For example, rare haplotypes that existed in East Africa via ILS before OOA may show a false high S* score if the rare haplotypes were included in OOA migrating population but were not included in the reference population in the S* analysis. Our obtained high S* scores of the H2 haplotype in the 17q21inv locus (Fig. 2) can be explained by no sampling of H2 Africans in this study, unlike Alves et al. (2015), and high colocalization of H1 and H2 within Eurasia, because of restricted recombination between H1 and H2 chromosomes.

### How to Treat “Locus”

We defined the “locus” as a LD region that was determined using the whole sample set of the 1000 Genomes Project. The concept of “locus” is operational and should be carefully treated in situations where genome-wide massive sequence data are available. We propose re-defining locus as a smaller LD region prior to phylogenetic analysis if specific haplogroups are disproportionally selected compared with others.

EHH analysis that focuses on an SNP can distinguish interesting haplogroups, such as those with unusual divergence, should be effective for identifying selection pressure only a specific allele. Moreover, EHH can display selection pressure differences within a genomic region as a density of bifurcation of haplotype lineages, as we showed. This is also effective for identifying differences in selection pressure among haplotypes that affect LD length. When the effect of heterogeneous selection is removed, the impact of variation in LD length by recombination alone becomes smaller, which facilitates phylogenetic analysis.

To estimate the actual effect of recombinants on phylogenetic analysis, we reviewed the position on our gene tree of haplotypes that corresponded to recombinants that were identified and removed from the phylogenetic analysis in the previous study on the HYAL locus, although our method that used imputation and phasing processes differed from the previous study (Ding et al. 2013). Contrary to expectation, we did not find that all of those haplotypes were located on long branches or separated from the other closely related haplotypes without recombination (Brown diamond in Fig. s5). Among the haplogroups that suggested recombination by forming a parallelogram in the phylogenetic network (i.e., haplogroups *CDE, IJK, NOT, ABS*), two haplogroups, *D* in *CDE* and *O* in *NOT*, contained haplotypes that corresponded to recombinants in the previous study (Fig. 2). Thus, this indicates that the effect on recombination is not serious, and a massive sequence data set can be analyzed without removing all recombinant candidates. In this situation, a phylogenetic network can reveal recombination events among haplotypes as large parallelograms (Fig. 2).

## Materials and Methods

### Data

We downloaded VCF and index (.tbi) files of chromosomes from the ftp site of the 1000 Genomes Project (ftp://ftp.1000genomes.ebi.ac.uk/;The 1000 Genomes Project Consortium 2015). The version of the VCF files was Phase 1 Version 3. Phasing for diploid autosomes was conducted in ShapeIt2. The file names include chromosome names and version information, “SHAPEIT2_integrated_phase1_v3.20101123.snps_indels_svs.genotypes.all.vcf.gz” for autosome, and “phase1_release_v3.20101123.snps_indels_svs.genotypes.all.vcf.gz” for the X chromosome.

The data contained 1,092 individuals from 14 populations (The 1000 Genomes Project Consortium 2012) (Table s2). The “American” samples used in the 1000 Genomes Project were determined to represent admixture of various North Americans that were more closely related to Africans than Native Americans (The 1000 Genomes Project Consortium 2012).

### Definition of Loci

We selected eight loci that contained candidate haplotypes for introgression according to the following criteria. First, the genomic regions were previously reported to include candidate haplotypes for which OOA cannot explain their divergence and/or geographic distribution pattern. Second, the sequence and genome coordinates of the candidate haplotype for introgression could be clearly detected based on the description in each previous paper.

#### Selection of focal haplotypes and SNPs

We manually inspected haplotype sequences reported by previous studies. Based on this inspection, we determined the most diverged haplotype as a haplotype of interest (focal haplotype) and an SNP site that specifically discriminated the focal haplotype (focal SNP) for each locus. When a LD block was not determined because of small minor allele count of the focal SNP, we selected focal haplotypes that represented exceptions to the OOA model in previous studies.

#### LD region determination

We calculated the *r^2^* values for all combinations of SNPs that existed within 200 kb in both directions of the ancient haplotype regions using data downloaded from the 1000 Genomes Project; for this, we used VCFtools with the --hap-r2 optional command (Danecek et al. 2011). We extracted SNPs that were closely associated (i.e., *r^2^* ≥ 0.8) with the focal SNP. We defined these LD regions as loci to be examined (Table 1).

In the application of our method to the 17q21inv locus, a genomic region with high LD (i.e., *r^2^* ≥ 0.8) was further examined to clarify the state of duplication within the LD region. The distribution of *r^2^* values within the LD region over 17q21 was divided into clusters according to a density-based clustering algorithm, Density Reachability And Connectivity Clustering (Ester et al. 1996), using fpc in the R library with the parameters ε=50000 and MinPts=50 (Hennig 2019). Based on the results of chr17:43654468–44369518, we eliminated the region that showed duplication and finally obtained chr17:43654468–44205122 as the region to be further analyzed. Consequently, the defined genomic region did not include the known and intensively focused SD that segregates sub-haplotype H2D in the H2 haplotype and H1D in the H1 haplotype (see Discussion).

### Data Validation

#### Data cleaning and re-genotyping

The obtained VCF files were trimmed according to the definition of loci. Because the VCF files contained regions with a short read-depth, we conducted a pilot investigation into whether base-calling of the VCF files might be improved by manual comparison with raw data (i.e., base-calling and quality values in BAM files) for five individuals per locus. This pilot study showed inconsistency between base-calling in VCF and quality in BAM, and insight into base-calling based on the quality values of read sequences.

Although the base and phase information of variant sites in VCF files that were congruent with raw data in BAM files were used for further analysis, we rewrote the VCF files to prioritize our observation of the raw data in the BAM file if there were inconsistencies among data sources.

The BAM files and index files for the target genomic regions were downloaded from the same ftp site for the VCF files mentioned earlier (ftp://ftp.1000genomes.ebi.ac.uk/;The 1000 Genomes Project Consortium 2015) using our in-house programs.

We eliminated information of reads described in the BAM files if there were more than two mismatches to the reference genome within 10 bases by replacing the base call to “N” and the quality value to “0”. We discarded PCR and optical duplicate reads.

We calculated quality values of the variant site described in the VCF files by combining quality values or read sequences obtained from the BAM files according to the algorithm in a program (ConstructAnalysis.py) developed by Brad Chapman (https://bitbucket.org/chapmanb/synbio/src/7b1b3a972b7e/SynBio/Sequencing/). See supplementary document for detailed information.

#### Imputation and re-phasing

Insertion and deletion (indel) sites in the VCF files, which were re-genotyped as necessary, were separated. Excluding those data, we divided the individual data in VCFs based on whether they contained unphased or missing sites. Then, imputation against missing base-call values was performed for each phased and unphased file followed by re-phasing using Beagle 3.3.2 (Browning and Browning 2007). Each result of the imputation was incorporated into the VCF file. Here, for heterozygous sites in Beagle output, we compared the phase of the heterozygous site with those of the neighboring three consecutive heterozygous sites. When the phases of these three sites did not match, the phase of the heterozygous site was recorded as ‘unknown phase.’ Otherwise, the phase assumed by Beagle was used for the new VCF file. To reduce such unknown information, we repeated imputation and re-phasing in Beagle using the obtained VCF file. Then, we confirmed that the renewed VCF contained fewer ‘missing’ bases and ‘unphased’ chromosomes (Table s4).

### Gene genealogy analysis of haplotypes

See supplementary document for information about preparation of sequences of Neanderthal, Denisovan, and chimpanzee.

#### NJ tree and bootstrapping

In the VCF files, we evaluated and corrected base-calling and phasing for the 1000 genomes, Altai Neanderthal, Denisovan, and chimpanzee were combined into one VCF file that included indel site information. Based on the variant information of the VCF files, nonredundant haplotype sequences were generated after removal of sequence data with 0.5% or more deleted sites in length.

With these haplotype sequence data for each locus, we constructed NJ trees (Saitou and Nei 1987) and added bootstrap values using a bash shell script, fasta2trebs.bsh, which automatically executes PHYLIP Dnadist for distance calculation between haplotypes under the F84 model of nucleotide substitution (Kishino and Hasegawa 1989; Felsenstein and Churchill 1996); PHYLIP Neighbor for NJ tree construction; and PHYLIP Seqboot for bootstrapping (using 500 iterations for each locus for this study) (Felsenstein 1989). This technique was previously published (Shimada and Nishida 2017).

We partially executed data validation of VCF files and NJ tree construction on the NIG supercomputer at ROIS National Institute of Genetics (Mashima et al. 2017).

#### Phylogenetic Network

Selection of Operational Taxonomic Units for the Phylogenetic Network:

To clarify the phylogenetic relationships among clusters shown on the NJ trees, we constructed a phylogenetic network with selected operational taxonomic units (OTUs) that represent each cluster of the NJ tree. Therefore, we developed an algorithm to select a small number (14–21 in this study) of OTUs and preferably maintain relationships among clusters without bias and arbitrariness. The algorithm includes the following two steps. First, OTUs were selected that comprehensively and homogeneously maintained their distances from each other. Second, OTUs with extraordinary distances from the root of each NJ tree were added to the OTUs selected in the first step. The in-house programs for these steps are available at an open repository (URL will be public after acceptance of the manuscript.) Briefly, the first program, tree_cluster.pl, determined the candidates of representative clusters and representative OTUs for the clusters. Then, the candidates of representative clusters were removed according to the distance to the neighboring candidate clusters. The removal steps were repeated until the number of candidate clusters reached the upper limit that was previously determined (parameter settings are provided in Table s1).

The second program, check_tree.pl, detected OTUs with extraordinary distance from the root of the NJ trees and added them to the OTUs that represented clusters in the first step. The haplotype sequences without indel sites of these selected OTUs and the two archaic hominins were saved as VCF files.

#### Construction of the Phylogenetic Network

The VCF file was transformed into an RDF file using an in-house perl script. Another VCF file with chimpanzee data was also used to check root position. We constructed a Reduced Median Network (Bandelt et al. 1995) with these RDF files using the free software Network 4.6 (http://fluxus-engineering.com). When too many parallelograms make it difficult to visualize and interpret, we adjusted the reduction threshold to reduce unnecessary median vectors and links by manual testing according to the user guide document (parameter settings are provided in Table s1).

### S* analysis

We conducted S* analysis that was originally devised for analyses with a small number of individuals, such as 20 (Plagnol and Wall 2006). Later, Vernot et al. (Vernot and Akey 2014; Vernot et al. 2016) extended this approach so it could be applied to a large number of individuals, and added a step to statistically quantify the matching between a candidate haplotype for introgression and an archaic haplotype. These previous S* calculation methods (Plagnol and Wall 2006; Vernot et al. 2016) were assumed to use non-phased data. Because we used phased data, we modified the original S* method (Plagnol and Wall 2006) so that it could be applied to phased haplotype data, including missing alleles, and was based on allele distance of a haplotype and not genotype distance on a non-African individual. To avoid obscuring the possibility of introgression from unknown archaic hominins by overemphasis of known archaic hominins, we simply displayed the haplotypes that deviated from the OOA model with classification by intensity of S* score and without quantification of matching to available archaic sequences.

We defined the reference population as the African populations LWK, YRI, and ASW (Table s2). We selected SNP sites with minor alleles observed in non-African haplotypes and allele frequencies in African less than 5% for use in S* calculation; this was to minimize the possibility of gene flow between non-African (target) and African (reference) populations. We separately calculated S* for three target populations (European, Asian, and American). To calculate S* of a haplotype, 100 haplotypes were randomly selected from the same target population.

#### Algorithm

We largely followed the sequence of steps for S* calculation described in previous studies (Plagnol and Wall 2006; Vernot and Akey 2014). However, we calculated S* by summing distances *d_h_*(*i.j*) between allele pairs of SNP sites *i* and *j* within haplotype *h*, and not genotype distance within a diploid individual. See supplementary documents for detailed algorithm.

#### Classification and display of *S\** results on the phylogenetic tree/network

To simplistically display S* intensities on phylogenetic trees and networks, we classified S* values into three classes in each locus. High and medium classes were defined as introgression grades I (*T2* ≤ S*) and II (*T1* ≤ S* < *T2*), respectively. We first determined *T1* and *T2* in dys44 and RRM2P4 loci by visual observation of distribution of S* values as *T1_dys44_*=60000, *T2_dys44_*=80000, *T1_RRM2P4_*=40000, and *T2_RRM2P4_*=53333, respectively. We chose these two loci because they slightly overlap with genic regions. Thresholds for other loci were calculated by assuming a linear relationship between the thresholds and number of SNPs, *N*, in these two loci, dys44 and RRM2P4, as follows:

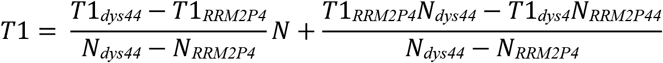

by assigning N_dys44_=313, N_RRM2P4_=209, and the above-mentioned values

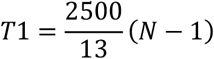

The second threshold T2 was defined as,

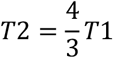

### EHH

We added ancestral allele information obtained from the UCSC genome browser to the VCF files, and we conducted data cleaning and re-imputation in Beagle 3.3.2 (Browning and Browning 2007). For the “rehh” R package (Gautier and Vitalis 2012), we generated two input files (i.e., for haplotypes and SNPs) from the VCF files with an inhouse perl program. We set a focal SNP site for EHH analysis as the same SNP site that was used to determine LD region. If multiple focal SNPs with perfect association (i.e., *r^2^*= 1) existed, a centrally located SNP was chosen. Because we applied the same criteria for choosing focal SNP sites for EHH analyses over all loci, the chosen SNP site was occasionally different from the “SNP for marker of introgressive haplotype” described in the original study, which happened at the HYAL locus; that is, rs116075629 was chosen instead of rs12488302 (Ding et al. 2013). We confirmed that the phylogenetic relationship of the HYAL haplotypes obtained in this study was equivalent to that in the original paper published by Ding et al. (2013), and this finding does not change the main argument regarding introgression from Neanderthals. We excluded haplotype data that contained many missing genotype sites by setting the min_perc_geno.hap=99.999 option of data2haplohh in the rehh program. EHH calculation results were represented in EHH plots. EHH regions were defined as genomic regions with EHH values ≥ 0.05 in both the ancestral and derived alleles of the focal SNPs. We also created a bifurcation graph within the regions with EHH values ≥ 0.2. In the case of MAF < 0.1, the obtained results were only used to identify genomic regions with stronger constraints by SNP density differences within the EHH region without comparing alleles to investigate selective sweep (Table s3), because haplotypes with rare SNP variants tend to have less haplotype variation, which elongates EHH in bifurcation graphs irrespective of selection.

## Acknowledgments

We are grateful to Michiyo Murase; Tsutomu Kanesashi, PhD; and Chiaki Inoue for help with the analyses, and to Jody Hey, PhD, for comments on an earlier version of this manuscript. We appreciate Takeru Akazawa, PhD, for leading the Replacement of Neanderthals by Modern Humans (RNMH) project, Kenichi Aoki, PhD, for leading the B01 research team of the RNMH, Ryosuke Kimura, PhD, for constructive discussion, and all the members of the RNMH for helpful comments and encouragements. We thank Mallory Eckstut, PhD, from Edanz Group (https://en-author-services.edanzgroup.com/) for editing a draft of this manuscript. This research was financially supported by the Ministry of Economy, Trade and Industry of Japan (METI), a Grant-in-Aid for Scientific Research on Innovative Areas (RNMH project 1201; Research in a proposed research area, Grant Number 23101506), JSPS the Grants-in-Aid for Scientific Research (C) (Grant Number 18K06452), and the Saito Gratitude Foundation to MKS.

## Abbreviations

(EHH): extended haplotype homozygosity
(ILS): incomplete lineage sorting
(indel): insertion and deletion
(LD): linkage disequilibrium
(MAF): minor allele frequency
(Mya): million years ago
(NJ): neighbor-joining
(OOA): out-of-Africa
(OTU): operational taxonomic unit
(*R.length*): genomic region length ratio of EHH to LD
(SDs): segmental duplications
(SNP): single nucleotide polymorphism

