## Supplemental materials for "Focusing on human haplotype diversity in numerous individual genomes demonstrates an evolutional feature of each locus"

### Data Validation

Calculation of quality values of the VCF files according to the algorithm in a program (ConstructAnalysis.py) developed by Brad Chapman:

The quality values of a variant site were calculated per site per sequencing platform (ILLUMINA, SOLID, or LS454) per sequencing project (low coverage or exome) by

$$Q = \sum_{k=1}^{n_f} \frac{Qf_k}{k} + \sum_{k=1}^{n_r} \frac{Qr_k}{k}$$

where $nf$ is number of reads with forward strands, $nr$ is number of reads with reverse strands, $Qf_k$ is the $k$-th largest Phred quality score in reads with forward strands, $Qr_k$ is the $k$-th largest Phred quality score in reads with reverse strands.

Based on our pilot investigation, we defined the following principles to genotype according to quality values in BAM files: $Qj$ ($Q1 \geq Q2 \geq Q3 > 0$) for bases with $j$-th highest quality among bases mapped at the position. First, a base with $Q < 40$ was not assumed to be true (i.e., change to '*N*' for the base). Second, if the ratio in quality values of the two bases mapped on a variant site was not greater than 5, $Q1/Q2 < 5$ ($Q1 \geq Q2$), the site was assumed to be heterozygous. By applying these principles, we rewrote the genotypes in VCF file using the following criteria:

genotypes re-evaluated for diploid chromosomes =

| | |
|---|---|
| heterozygous ($Q1/Q2$) | if $Q2 \geq 40$, $Q1/Q2 < 5$, and $Q2 - Q3 > 5$, |
| heterozygous ($Q1/N$) | if $Q2 \geq 40$, $Q1/Q2 < 5$, and $Q2 - Q3 \leq 5$, |
| | or if $Q1 \geq 40$, $Q1/Q2 < 5$, and $Q2 < 40$, |
| homozygous of $Q1$ bases ($Q1/Q1$) | if $Q1 \geq 40$ and $Q1/Q2 \geq 5$, |
| homozygous of "$N$" ($N/N$) | if $Q1 < 40$ |

genotypes re-evaluated for haploid chromosomes =

| | |
|---|---|
| $Q1$ | if $Q1 \geq 40$ and $Q1 - Q2 > 5$ |
| $N$ | otherwise |

### Gene genealogy analysis of haplotypes

Preparation of Neanderthal and Denisovan sequences:

VCF files for chromosomes of an Altai Neanderthal and Denisovan were downloaded from the Max Planck Institute for Evolutionary Anthropology website (http://cdna.eva.mpg.de/neandertal/altai/AltaiNeandertal/VCF/, http://cdna.eva.mpg.de/denisova/VCF/hg19_1000g/).

After extracting target genomic regions from the chromosomal VCF files, haplotype sequences were determined from the downloaded ancient hominin sequences as follows: (1) bases at sites with "LowQual" in the FILTER field were replaced with alleles of human reference sequences, (2) cases other than those described in (1) for which we determined haplotype sequence from diploid data by selecting an allele with more read counts for single base substitution sites and selecting reference alleles for indel sites.

Preparation of Chimpanzee sequence data as an outgroup:
We obtained chimpanzee sequences for the target genomic regions from the UCSC Table browser (http://genome.ucsc.edu/cgi-bin/hgTables) by selecting the following options: clade: Mammal, genome: Human, assembly: Feb. 2009 (GRCh37/hg19), group: Comparative Genomics, track: Conservation, table: Multize Align (mulitz100way), output format: MAF – multiple alignment format.

### S* analysis

Algorithm of S* calculation by summing distances $d_h(i,j)$ between allele pairs of SNP sites $i$ and $j$ within haplotype $h$:
Calculation of phased total distance over all haplotypes, *distance*$(i,j)$:
We modified the calculation of genotype distance $d(i,j)$ to use phased data as follows:

$$distance(i,j) = \sum_h d_h\,(i,j)$$

where $d_h(i,j)$ is coded as 1 for a combination of major and minor alleles and 0 for other combinations. According to a previous study (Plagnol, V and Wall, JD 2006), we scored $S_0\,(i,j)$ for SNP pairs $(i, j)$ within a haplotype as follows:

$$S_0(i,j) = \begin{cases} bp(i,j) + 5000 & (distance(i,j) = 0) \\ -10000 & (1 \leqq distance(i,j) \leqq 5) \\ -\infty & (5 < distance(i,j)) \end{cases}$$

where $bp(i,j)$ is the distance between genomic positions of the two SNPs $(i, j)$.
When missing data were included, we applied the criteria used in the previous study (Plagnol, V and Wall, JD 2006). Briefly, a target population contained no more than two chromosomes with missing calls at an SNP associated with a minor allele of the other SNP in an SNP pair. Moreover, only one such chromosome should exist when the MAF is 2 in the population. If the criteria were met for an SNP pair $(i, j)$, then we counted $S_0(i,j) = bp(i,j) + 5000$ for $distance(i,j) = 0$; otherwise, it was set as $S_0(i,j) = 0$.
We used so-called dynamic programming that reduced computation quantity during search of the maximum value of $S\,(J)$ according to previous studies (Plagnol, V and Wall, JD 2006; Vernot, B and Akey, JM 2014) as follows:

$$S_x = \max S(J_x) = \max_{1 \leq y \leq x-10} \{\max(S_y, 0) + S_0(y,x)\}$$

where $1 \leqq y < x$ and $bp(y, x) \geqq 10$, $J$ is a list of SNPs ordered by genomic position, and $J_x$. is the last SNP in $J$. To find $S*$, we calculated $S_x$ and $S_y$ for all possible $x$, $y$ pairs in ascending order of $x$ as follows:

$$S^* = \max_{1<x\leq n} S_x$$

### Investigation of removed haplotype in the previous study of HYAL locus

To evaluate impact of recombination to phylogenetic analysis, we focused on removed haplotypes in the previous study (Ding et al. 2013) as recombinants between introgressed and non-introgressed segments. In the previous study, derived alleles of 26 specific SNPs (Type-2 SNPs) were considered introgressive haplotypes, and 19 haplotypes bearing both derived (introgressive) and ancestral (non-introgressive) alleles among the 26 Type-2 SNPs were considered recombinants and were removed. In the current study, haplotypes of the HYAL locus were determined in a different way (i.e., in locus definition and phase determination) from the previous study, and recombinant candidates were not removed. Then, correspondence was determined between recombinants from the previous study and the current study.

We investigated the sample ID of individuals that have recombinant haplotypes in the previous study. Among the haplotypes obtained from these samples, we compared the obtained haplotypes in the two studies and determined correspondence between them.

Haplotypes from the current study are shown below, and corresponding haplotypes from previous studies are indicated in parentheses.

NA18572-1 (NA18572 b)
NA18609-2 (NA18609 a)
NA18641-1 (NA18641 b)
HG00501-1 (HG00501 a)
HG00590-1 (HG00590 b)
HG00634-2 (HG00634 b)
HG00662-2 (HG00662 b)
HG00692-1 (HG00692 a)
NA18950-1 (NA18950 a)
NA18950-2 (NA18950 b)
NA18968-1 (NA18968 b)
NA18977-1 (NA18977 a)
NA18995-1 (NA18995 b)
NA19735-2 (NA19735 a)
HG01271-2 (HG01271 b)

Haplotypes incomplete correspondence with candidate in previous study
HG00651-1 (HG00651 a)
NA18945-2 (NA18945 a)
HG00701-2 (HG00701 a)
HG00436-2 (HG00636 a, possibly typo of HG00436 a)

Then, we investigate alleles of all SNPs of the haplotypes in current study and classified whether the haplotypes were corresponding to haplotype as recombinants in the previous study (closed diamond in Fig. s5), or not (open circle in Fig. s5). These haplotypes were subdivided from the view of recombination between ancestral and derived genomic segments according to alleles of the Type-2 (Fig. s5).

### Supplementary Figure s1:

NJ trees for haplotypes of modern humans and archaic hominins (Altai Neanderthals and Denisovans). (a) Xp11hs, (b) dys44, (c) RRM2P4, (d) 17q21inv, (e) STAT2. Those of three other loci (MCPH1, OAS, and HYAL) are depicted in Figure 1. See Figure 1 for detailed information about the style of NJ trees.

### Supplementary Figure s2:

Distribution and maximum of S* values with two determined thresholds. Positive S* values were classified into four intervals whose endpoints are written in E-notation to avoid superscript use. Specifically, *m*e+*n* indicates a value of $m \times 10^n$. See Materials and Methods for the calculation formula of S* and determination of the two thresholds.

### Supplementary Figure s3:

Sequence alignment of SNP sites within the genomic region for EHH bifurcation graph (EHH values $\geq 0.2$) in the MCPH1 locus: 20 haplotypes of 1000 genomes, and Denisovan and Neanderthal haplotypes that were selected for phylogenetic network construction are displayed, although EHH analysis was conducted with all sequence data of 1000 genomes. Yellow background indicates extended haplotype regions carrying the ancestral allele (G) at the focal SNP (rs930557). Bold SNPs split these haplotypes by polymorphisms that correspond to bifurcations in the bifurcation graph of the ancestral allele, which is shown by red lines in Figure 5C.

### Supplementary Figure s4:

EHH plots and bifurcation graphs. (A) Xp11hs, (B) dys44, (C) RRM2P4, (D) 17q21inv, (E) STAT2, (F) OAS, (G) HYAL. See Figure 5 for the MCPH1 locus and detailed information about these graphs.

### Supplementary Figure s5:

Correspondence was determined between recombinants from a previous study (Ding et al. 2013) and the current phylogenetic analysis of the HYAL locus. Based on all SNPs of the HYAL loci in current study, haplotypes from the samples that have recombinant haplotypes in the previous study were classified whether corresponding to haplotype as recombinants in the previous study (closed diamond), or not (open circle). Colors indicate status in haplotype of current study about recombination between derived (or introgressed from Neanderthal) and ancestral (non-introgressed) genomic segments based on 26 SNPs used to infer introgression in the previous study: such recombination was observed (Brown), no recombination was observed but derived alleles were observed (Green), and no recombination was observed but derived alleles were observed (Red). See supplementary document for detailed information.

Table s1. Summary of comparison between phylogenetic trees and networks

| # | Locus | Tree vs. Network | BS value around root | Size and frequency of reticulation (parallelogram) in phylogenetic network | Derived allele distribution of focal SNP in haplotype phylogeny | N-D relation(1) | NJ-tree topology pattern(2) | Parameters Representative OTU (tree_cluster.pl) --size | --interval | --neighbo | --reps | Network4.6 Reduction threshold |
|---|---|---|---|---|---|---|---|---|---|---|---|---|
| I | Xp11hs | Partly different, e.g., clusters *L* ,*M* , and *N* | High | Edges to the most diverged cluster form series of many thin and long parallelograms | Collectively distributed in the outmost cluster including haplotypes *A* and *B* | + | ((A,C),(A,((N,D),(A,(A,(A,C)))))) | 20 | 0.0004 | 0.0002 | 1 | 5 |
| II | dys44 | Partly different, e.g., clusters *F* , and *L* | High | Shows many parallelograms offering various split pattern for many clusters | Four SNPs including the focal SNP constitute a common dimension among haplotypes *B* , *C* , Neanderthal, and Denisova in phylogenetic network suggesting recombination; the singleton edge to haplotype *T* contains two of the four SNPs (focal SNP and neighboring one), which makes isolated | - | (D,(E,(C,(N,C))),(E,(C,(A,(C,(A,E))) ))) | 20 | 0.001 | 0.0004 | 1 | 4 |
| III | RRM2P4 | Concordant, except cluster *P* | Low | Large parallelogram linked to clusters *G* to *I* | Separated haplotype C with 2-edges from other haplotypes of derived allele | - | ((A,(E,(C,(E,N)))),(C,(C,(A,E)))) | 10 | 0.001 | 0.0005 | 1 | 3 |
| IV | MCPH1 | Partly different, e.g., clusters *D* , *N* , and *R* | High | Multiple wide parallelograms representing recombinations, Small parallelograms offering various split pattern for many | Continuous distribution on the NJ tree but not on the phylogenetic network, which is brought by difference of algorithms that can correct with weighting based on synapomorphic SNPs (See discussion) | ++ | (A,((A,E),((A,(E,(N,D))),(C,(A,C))))) | 10 | 0.001 | 0.0012 | 1 | 4 |
| V | 17q21inv | Concordant | High | Large parallelograms linking to clusters A to D distantly located from root, which indicates recombination within a haplotype family with identical chromosome | Derived alleles are found in H1 group (clusters *A* to *D* ) that clearly divided by inversion; sub-Saharan African *H2'* cluster found in Alves et al. (2015) was not reproduced perhaps because our method did not examine sufficient number of Africans | ++ | (C,(C,((A,(N,D)),E))) | 5 | 0.0005 | 0.0015 | 20 | 2 |
| VI | STAT2 | Concordant | High | Wide parallelogram linked to clusters *A* to *E* suggesting recombination among them | Separated haplotype of Neanderthal from other haplotypes (*B* to *E* ) bearing derived allele can be explained by interupted clustering of haplotype *A* that is recombinant at focal SNP region | - | (D,((C,N),(E,(A,C)))) | 4 | 0.0001 | 0.0002 | 2 | 2 |
| VII | OAS | Concordant | High | Elongated parallelograms around root split two major clusters | Collectively distributed in haplotypes *H* to *Q* as well as *F* that are suggeste to be a recombinant at focal | - | ((C,(E,N)),(E,(C,(A,D)))) | 10 | 0.001 | 0.0002 | 1 | 2 |
| VIII | HYAL | Partly different, e.g., culsters *F* and *P* | High | Small parallelograms located around center of network offering various split patterns | Only the outmost cluster including haplotype *A* carries derived allele (blue); rs12488302-T presented by Ding et al (2013) located on parallel edges that formed clumped distribution of haplotypes *C,D,E,P* and Altai Neanderthal (brown) | - | (A,((D,(E,N)),(A,(C,(A,C))))) | 10 | 0.0001 | 0.00016 | 1 | 2 |

(1) Clustering of Neanderthals (N) and Denisovans (D), (-) not observed; (+) observed with 75% < BS < 99%; (++) observed with BS more than or equql to 99%.

(2) Framework of NJ tree tolopoly in Newick format; (A) African; (E) Eurasian; (C) Cosmoplitan; (N) Neanderthal; (D) Denisovan clusters. Red indicates typical OOA pattern.

Table s2. Samples and populations obtained from 1000 Genomes (phase 1 version 3) used in this study.

| Super Pop. Code | # in Continent | Population Code | # in Population | #Male | #Female |
|---|---|---|---|---|---|
| ASN | 286 | CHB | 97 | 44 | 53 |
| | | JPT | 89 | 50 | 39 |
| | | CHS | 100 | 50 | 50 |
| EUR | 379 | CEU | 85 | 45 | 40 |
| | | TSI | 98 | 50 | 48 |
| | | GBR | 89 | 41 | 48 |
| | | FIN | 93 | 35 | 58 |
| | | IBS | 14 | 7 | 7 |
| AMR | 181 | MXL | 66 | 31 | 35 |
| | | PUR | 55 | 28 | 27 |
| | | CLM | 60 | 29 | 31 |
| AFR | 246 | LWK | 97 | 48 | 49 |
| | | YRI | 88 | 43 | 45 |
| | | ASW | 61 | 24 | 37 |

Han Chinese in Beijing, China (CHB); Japanese in Tokyo, Japan (JPT); Han Chinese South, China (CHS); Utah residents with ancestry from Northern and Western Europe, US (CEU); Toscani in Italia (TSI); British from England and Scotland, UK (GBR); Finnish in Finland (FIN); Iberian populations in Spain (IBS); Mexican ancestry in Los Angeles, California, USA (MXL); Puerto Rican in Puerto Rico (PUR); Colombian in Medellin, Colombia (CLM); Luhya in Webuye, Kenya (LWK); Yoruba in Ibadan, Nigeria (YRI); African ancestry in Southwest USA (ASW); Asian (ASN); European (EUR); American (AMR); African (AFR). Detailed information can be found in IGSR: The International Genome Sample Resource (https://www.internationalgenome.org/faq/which-populations-are-part-your-study).

Table s3. Comparison of genomic region lengths between LD and EHH analyses

| # | Locus | Defined haplotype length [bp] (1) | Observed high EHH region (EHH≧0.2) [bp] | Length ratio (EHH/LD) (2) | SNP # ratio (EHH/LD) (2) | SNP density ratio (EHH/LD) | Estimated extent of recombination | MAF＞0.1 | LD-stretch ratio between alleles at EHH=0.5 | Difference b/w alleles in concurrent bifurcation | EHH summary |
|---|---|---|---|---|---|---|---|---|---|---|---|
| I | Xp11hs | 83,110 | 24,392 | 0.293 | 0.388 | 1.323 | +/- | - | n/a | n/a | Little difference in region length between LD and EHH suggesting stable genomic region; fewer branching points in upper region suggest functional constraint of overlapping with promoter of SHROOM4 gene (chrX:50,334,643-50,557,044) |
| II | dys44 | 35,162 | 11,048 | 0.314 | 0.269 | 0.856 | +/- | - | n/a | n/a | Little difference in region length between LD and EHH suggesting little impact by allele-specific selection |
| III | RRM2P4 | 23,198 | 7,433 | 0.320 | 0.312 | 0.973 | +/- | - | n/a | n/a | Comparatively larger *R.length* ; too small MAF compare to that of major |
| IV | MCPH1 | 67,083 | 1,927 | 0.029 | 0.028 | 0.983 | ++ | + | 0.05 | ++ | Major allele derived at possibly Eurasia strongly maintained longer EHH region than ancestral one, which suggests recombination rate differences among haplotypes that makes systematic error in phylogenetic analyses |
| V | 17q21inv | 550,655 | 2,737 | 0.005 | 0.006 | 1.191 | ++ | + | 0.71 | +/- | The smallest *R.length* (0.5%) among the loci, which implies substantial isolation between *H1* and *H2* haplotype families but existence of recombination within each of them; selective sweep on *H2* lineage (Stefansson et al 2005) was not confirmed |
| VI | STAT2 | 130,476 | 18,136 | 0.139 | 0.134 | 0.965 | + | - | n/a | n/a | Too small MAF to compare with major allele |
| VII | OAS | 30,900 | 14,277 | 0.462 | 0.461 | 0.997 | +/- | + | 0.71 | +/- | Skewed distribution of branching points suggest selection-pressure difference at OAS1 gene (chr12:113,344,739-113,357,712) in the locus |
| VIII | HYAL | 176,931 | 11,820 | 0.067 | 0.090 | 1.345 | + | - | n/a | n/a | Skewed distribution of branching points suggests selection-pressure difference at HYAL3 gene (chr3:50,330,259-50,336,899) in the locus; showed longer EHH at allele of the most diverged haplogroup although too small MAF to compare with major allele |

(1) LD region, $r^2 \geq 0.8$

(2) Ratio ≤ 0.03, 0.03 < ratio < 0.2, and 0.2 ≤ ratio are depicted by yellow, yellowish green, and green, respectively

Table s4. Results of repeated imputation using Xp11hs VCF files

| Step of Imputation | # chromosomes | | | | # [sites × individuals] | | |
|---|---|---|---|---|---|---|---|
| | phased | | unphased | Total | Determined | missing or unphased | Total |
| | without missing | with missing | - | | | | |
| Original | 0 | 667 | 992 | 1659 | 612644 | 146296 | 758940 |
| 1st | 1512 | 119 | 28 | 1659 | 758719 | 221 | 758940 |
| 2nd | 1656 | 1 | 2 | 1659 | 758938 | 2 | 758940 |
| 3rd | 1659 | 0 | 0 | 1659 | 758940 | 0 | 758940 |

Table s5. Difference in consequences expected from population models

| | ILS of Ancestral polymorphism | Introgression via hybridization | References |
|---|---|---|---|
| Time since population/species division | More frequent in shorter divergence time | Independent of divergence time, Hard to recognize in shorter period | - |
| Expected minimum sequence divergence between from two populations/species | Larger | Smaller | Joly, et al. 2009 |
| Geographic distribution of a specific rare allele | Regionally scattered | Regionally intensive | Green, et al. 2010; Zhou, et al. 2017 |

Table s6. Sample origins of 25 haplotypes (sample ID + phase) in diverged clusters of the HYAL locus

| Cluster | Haplotype | Pop. Code |
|---|---|---|
| A | NA20341-1 | ASW |
| A | NA19904-1 | ASW |
| A | NA20296-2 | ASW |
| A | NA20289-2 | ASW |
| A | HG01060-2 | PUR |
| A | NA20356-2 | ASW |
| A | NA19469-2 | LWK |
| A | NA18489-1 | YRI |
| B | NA19399-1 | LWK |
| B | NA18874-2 | YRI |
| B | NA19359-1 | LWK |
| B | NA19381-1 | LWK |
| B | NA19107-2 | YRI |
| B | NA20281-2 | ASW |
| B | NA20276-2 | ASW |
| B | NA19428-1 | LWK |
| B | NA18861-2 | YRI |
| B | NA19257-1 | YRI |
| B | NA19444-2 | LWK |
| B | NA19451-1 | LWK |
| B | NA19036-2 | LWK |
| B | NA19434-1 | LWK |
| B | NA18909-2 | YRI |
| B | NA19309-1 | LWK |
| B | NA19200-2 | YRI |

Table s7. Sample origins of five haplotypes (ID + phase) in diverged clusters of the Xp11hs locus

| Cluster | haplotype | Pop. Code |
|---|---|---|
| A | NA19472-2 | LWK |
| B | NA20334-1 | ASW |
| B | NA18871-1 | YRI |
| B | NA19352-1 | LWK |
| B | NA20348-1 | ASW |

Table s8. Information on samples in diverged clusters in the HYAL and Xp11hs loci

| # in HYAL | # in XP11hs | Pop. Code | Super Pop. Code | Locality |
|---|---|---|---|---|
| 10 | 2 | LWK | AFR | Kenya, East Africa |
| 7 | 1 | YRI | AFR | Nigeria, West Africa |
| 7 | 2 | ASW | AFR | SW US (detail unknown) |
| 1 | 0 | PUR | AMR | Puerto Rico |

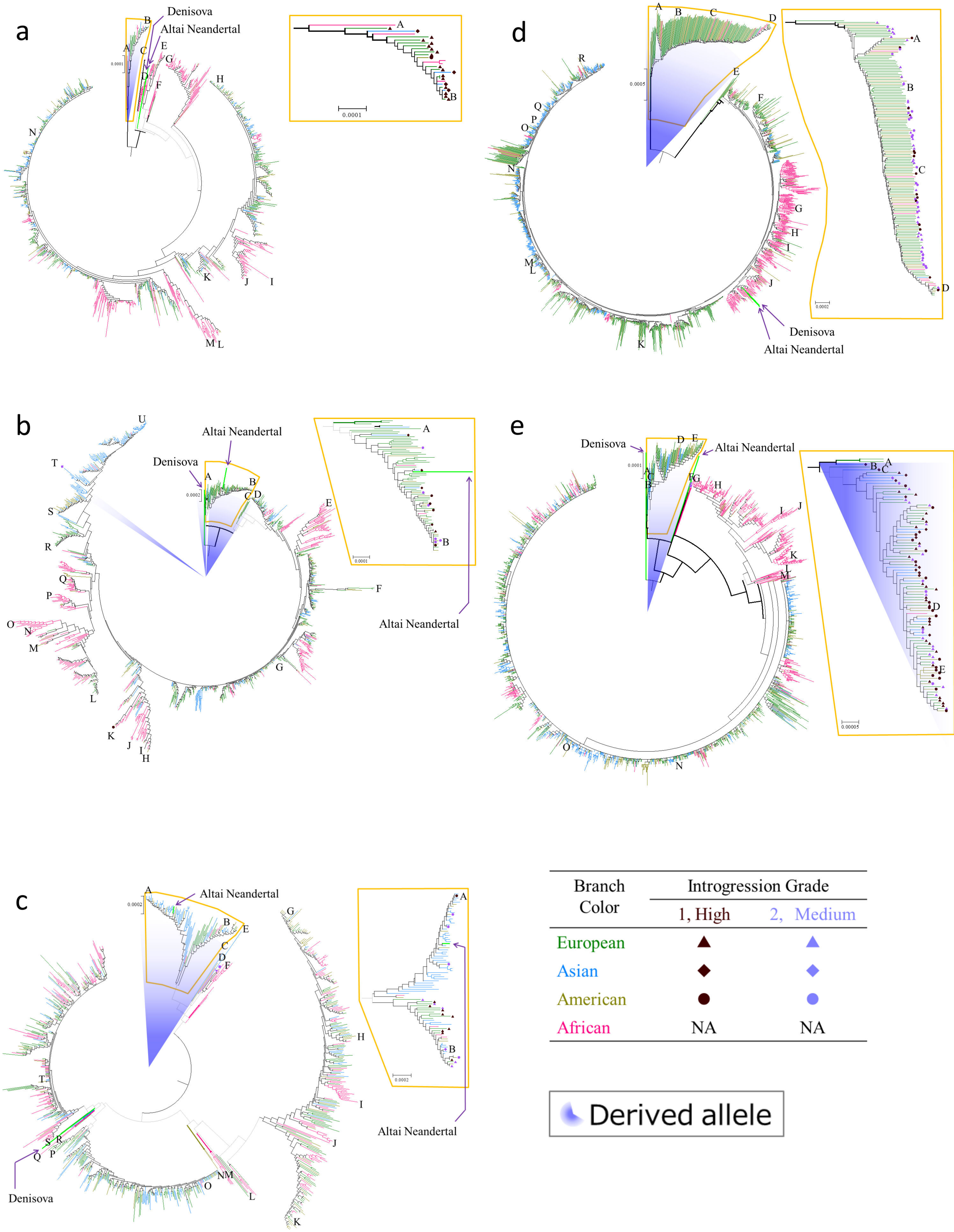
a
Denisova
Altai Neandertal
0.0001
b
Altai Neandertal
Denisova
0.0002
0.0001
Altai Neandertal
c
Altai Neandertal
0.0002
0.0002
Altai Neandertal
Denisova
d
0.0005
0.0002
Denisova
Altai Neandertal
e
Denisova
Altai Neandertal
0.0001
0.00005
Branch Color
Introgression Grade
1, High
2, Medium
European
Asian
American
African
NA
NA
Derived allele

Xp11hs

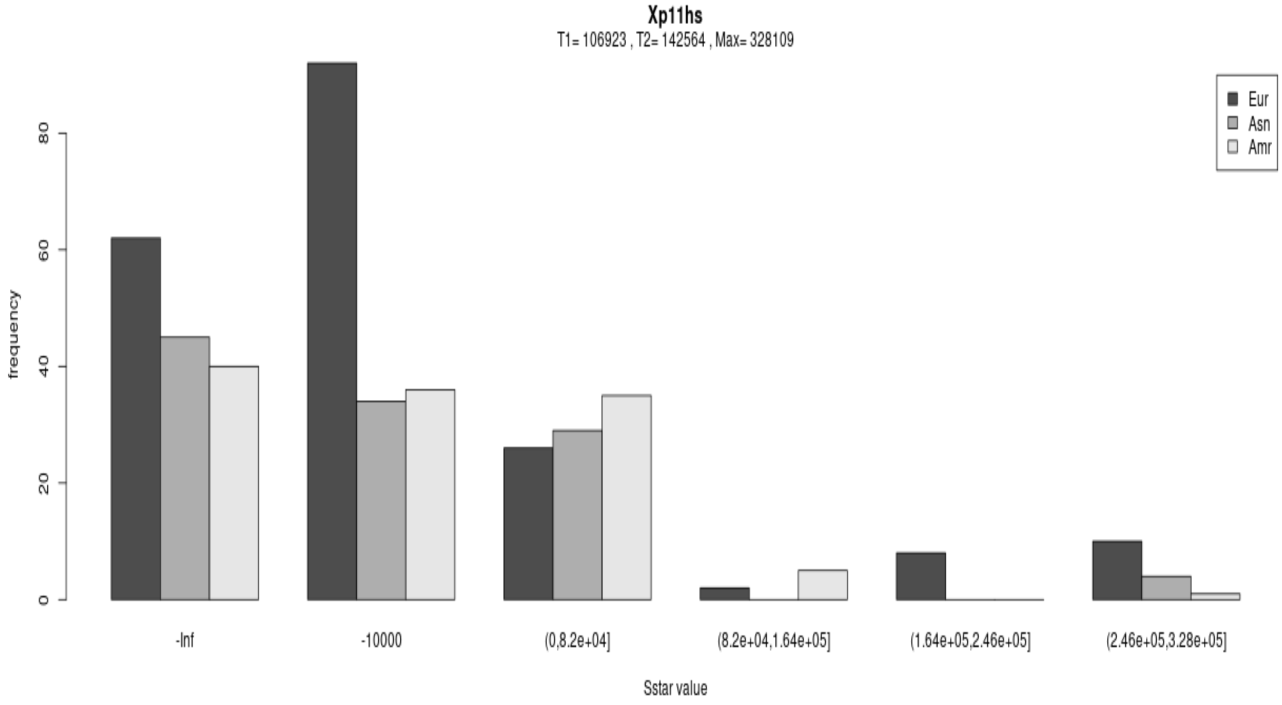


dys44

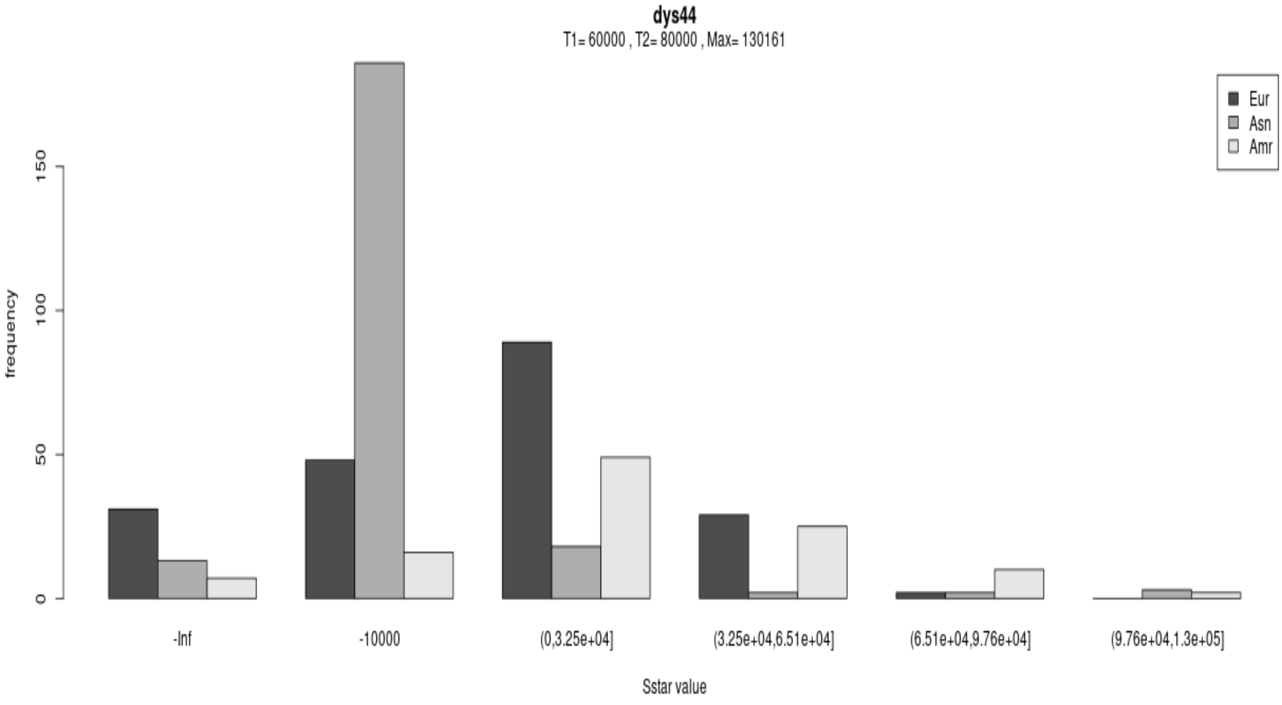

RRM2P4

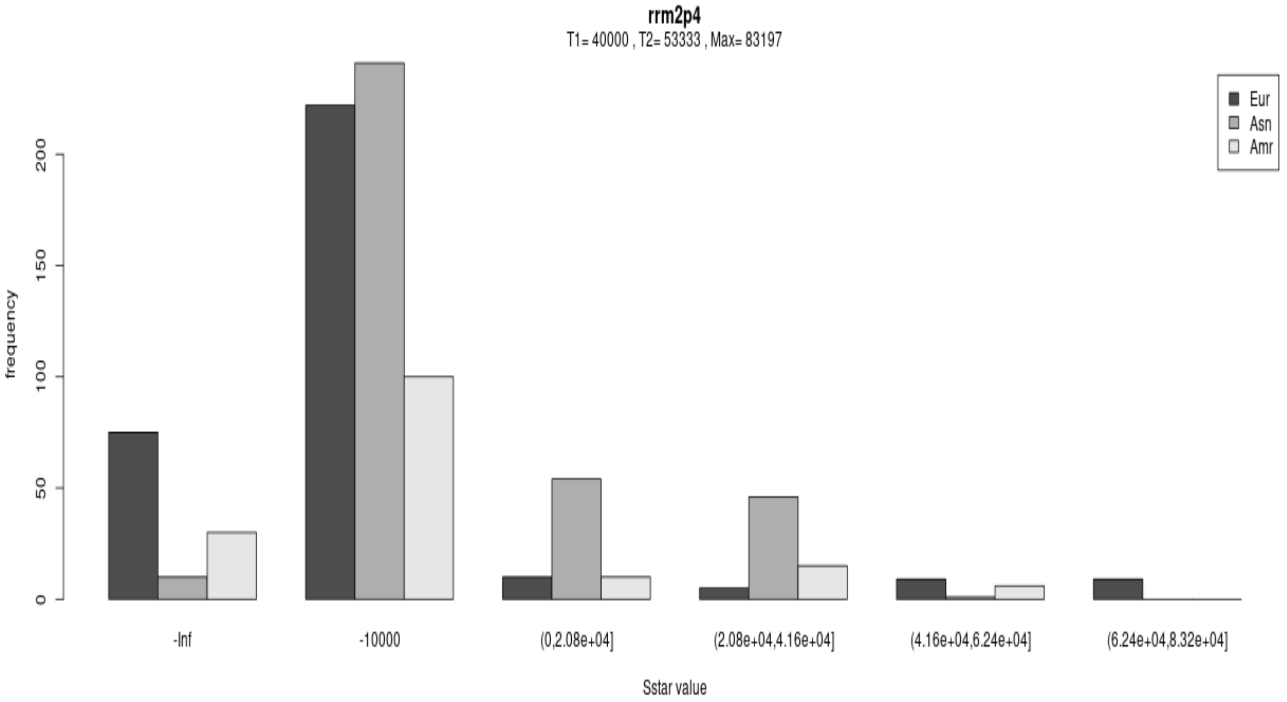


MCPH1

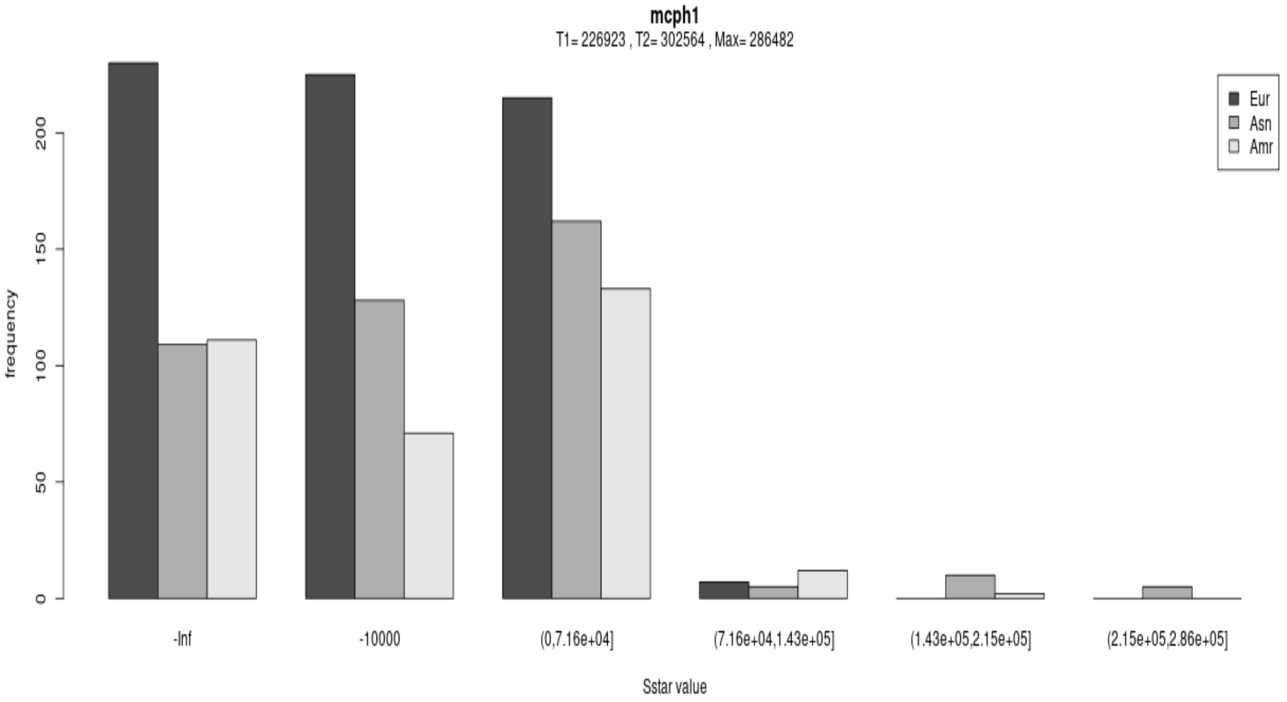

17q21inv

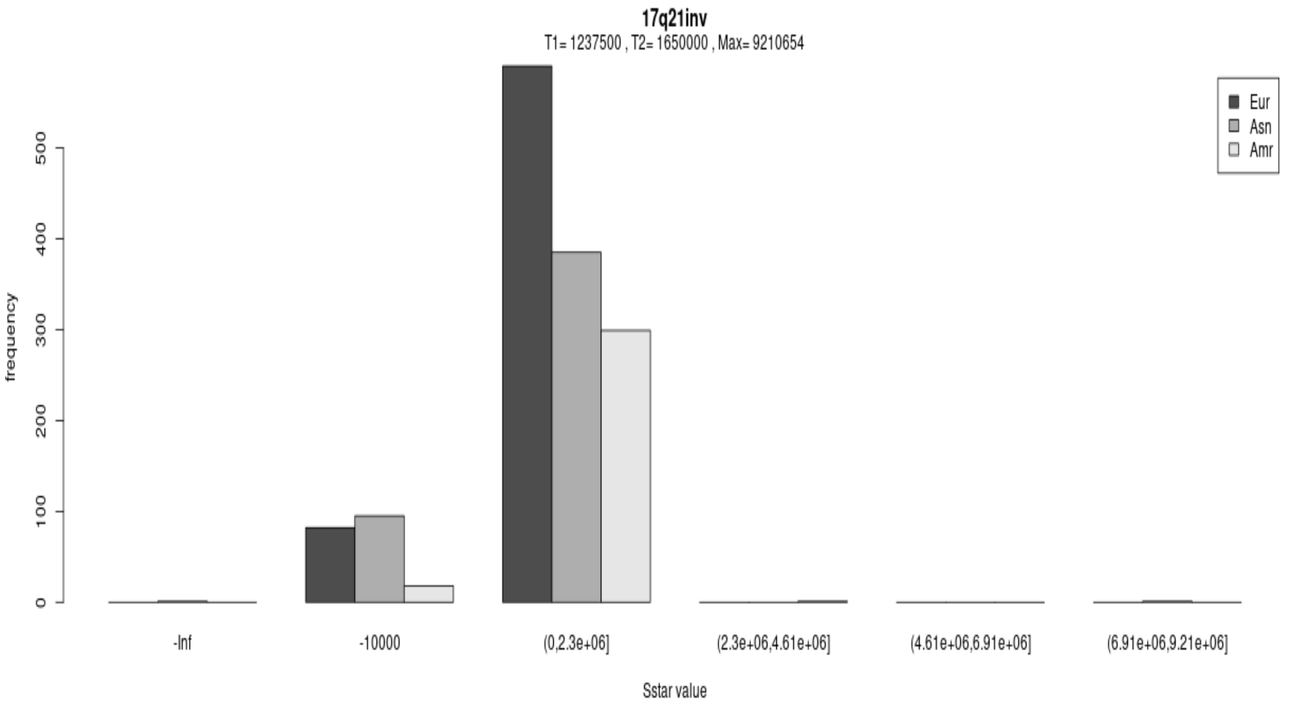


STAT2

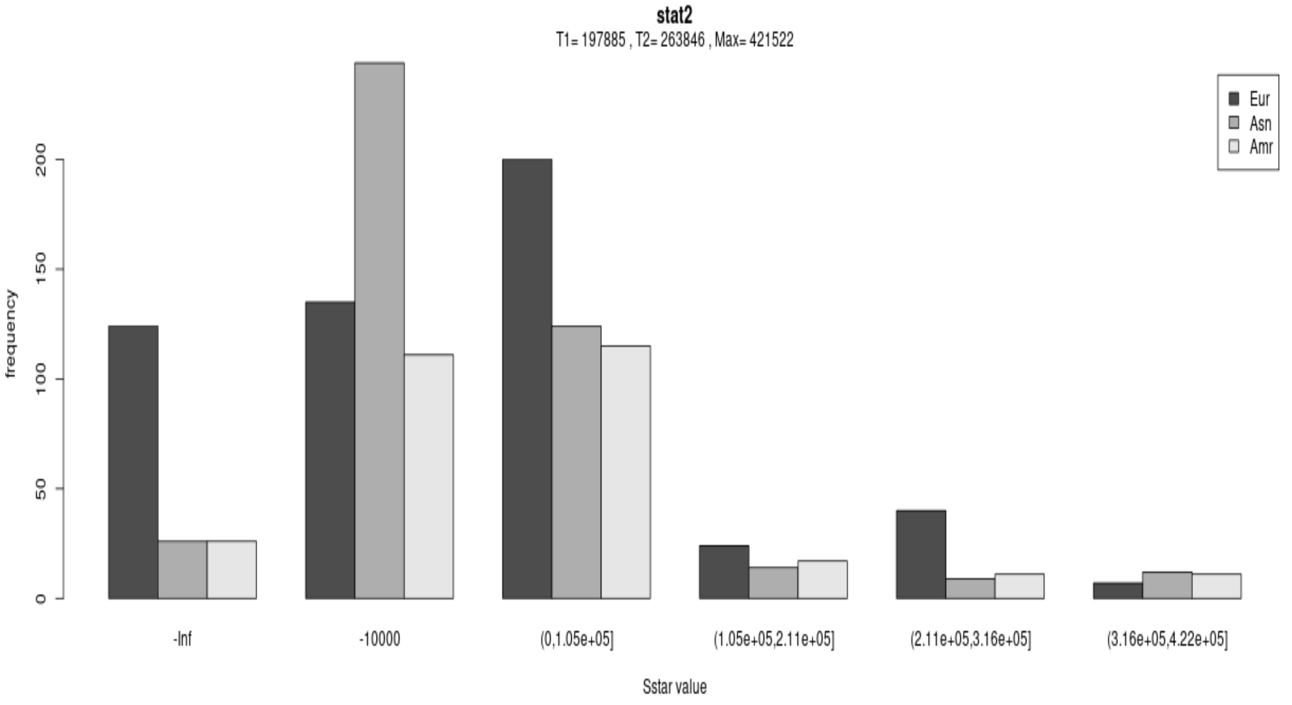

OAS

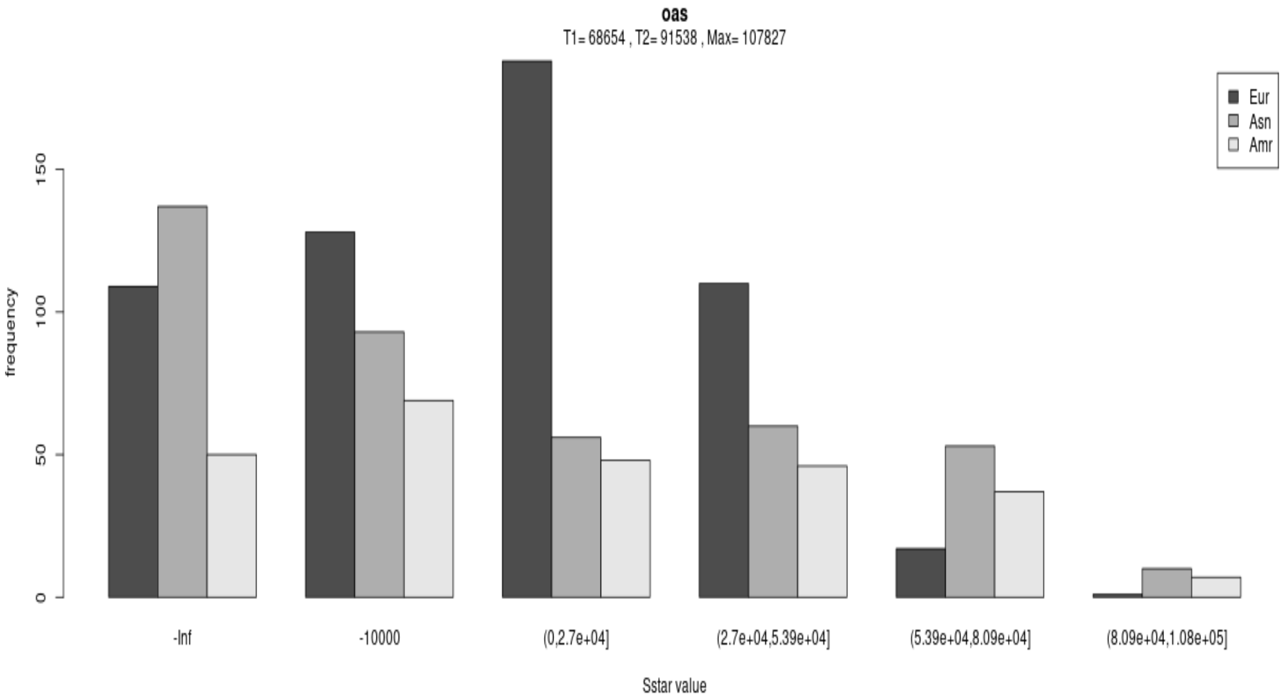
oas
T1= 68654 , T2= 91538 , Max= 107827
Eur
Asn
Amr
frequency
0
50
100
150
-Inf
-10000
(0,2.7e+04]
(2.7e+04,5.39e+04]
(5.39e+04,8.09e+04]
(8.09e+04,1.08e+05]
Sstar value

HYAL

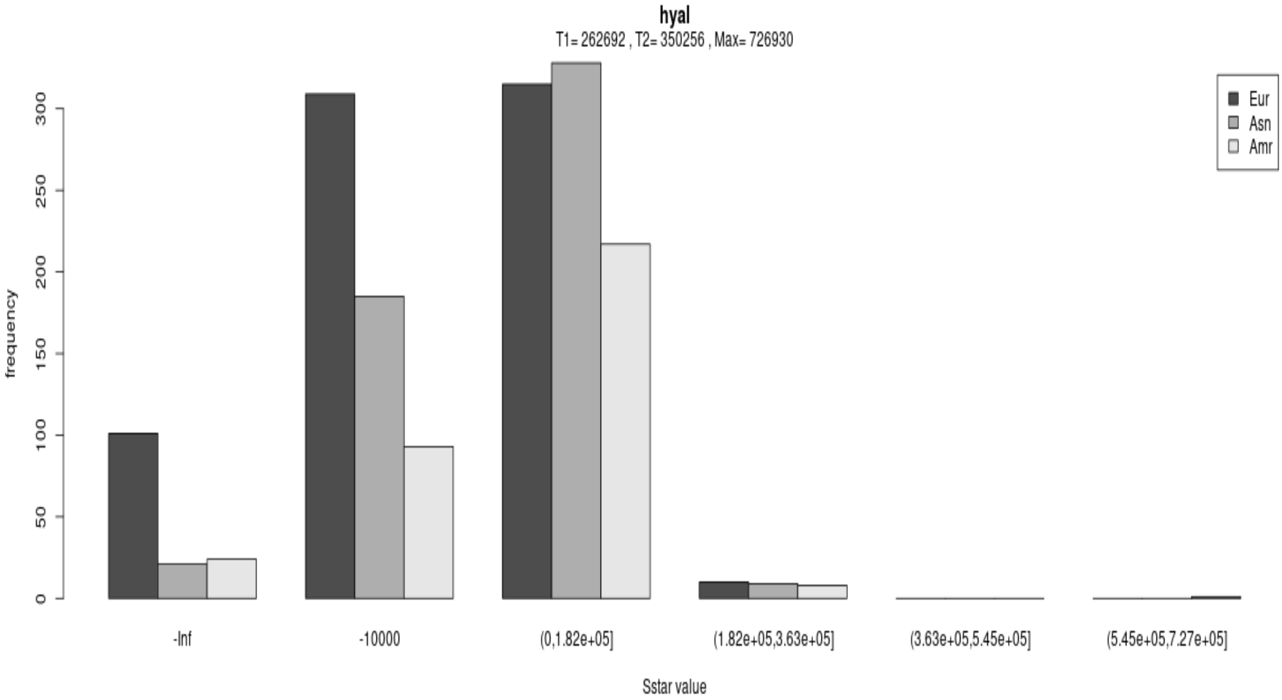
hyal
T1= 262692 , T2= 350256 , Max= 726930
Eur
Asn
Amr
frequency
0
50
100
150
200
250
300
-Inf
-10000
(0,1.82e+05]
(1.82e+05,3.63e+05]
(3.63e+05,5.45e+05]
(5.45e+05,7.27e+05]
Sstar value

| rsID | rs2440416 | rs189522330 | rs75666535 | rs182226349 | **rs11137028** | rs185966191 | rs62496812 | rs142720060 | rs147324953 | rs180825999 | rs201947779 | rs34121009 | rs141024274 | rs149813931 | rs201231900 | **rs2083914** | rs200406468 | **rs930557** | rs185613641 | rs115088000 | rs199750676 | rs145820898 | rs148526209 | rs190026099 | rs201084851 | rs200513127 | rs2515569 | rs142858644 | rs199724219 | rs77959215 | rs201039834 | **rs2920676** | rs185736569 | rs192003514 | rs183880522 | **rs146586991** | rs188726432 | rs141218500 | **rs35551093** | **rs41313954** | rs17076894 | rs146744659 | rs192777656 | rs182472674 | rs187245785 | rs140402469 |
|---|---|---|---|---|---|---|---|---|---|---|---|---|---|---|---|---|---|---|---|---|---|---|---|---|---|---|---|---|---|---|---|---|---|---|---|---|---|---|---|---|---|---|---|---|---|---|
| chr8(hg19) | 6301472 | 6301481 | 6301519 | 6301540 | **6301546** | 6301603 | 6301738 | 6301859 | 6301898 | 6301929 | 6301980 | 6302033 | 6302062 | 6302102 | 6302110 | **6302154** | 6302178 | **6302183** | 6302221 | 6302232 | 6302279 | 6302295 | 6302304 | 6302343 | 6302344 | 6302391 | 6302418 | 6302479 | 6302487 | 6302592 | 6302612 | **6302671** | 6302686 | 6302701 | 6302723 | **6302738** | 6302907 | 6302959 | **6302962** | **6302971** | 6302981 | 6303028 | 6303050 | 6303140 | 6303265 | 6303398 |
| HG00114R | G | A | T | T | G | G | T | A | C | G | A | A | T | A | G | G | C | C | T | A | C | G | T | C | G | C | G | T | A | A | G | C | C | A | G | G | C | C | C | C | A | T | C | A | C | G |
| HG00422R | G | A | T | T | G | G | T | A | C | G | A | A | T | A | G | G | C | G | T | A | C | G | T | C | G | C | G | T | A | A | G | C | C | A | G | G | C | C | C | C | A | T | C | A | C | G |
| HG00740R | C | A | T | T | G | G | T | A | C | G | A | A | T | A | G | G | C | G | T | A | C | G | T | C | G | C | G | T | A | A | G | C | C | A | G | G | C | C | C | C | A | T | C | A | C | G |
| NA18522R | C | A | T | T | G | G | T | A | C | G | A | A | T | A | G | G | C | G | T | A | C | G | T | C | G | C | G | T | A | A | G | C | C | A | G | G | C | C | C | C | A | T | C | A | C | G |
| NA18868R | C | A | T | T | A | G | T | A | C | G | A | A | T | A | G | G | C | G | T | A | C | G | T | C | G | C | G | T | A | A | G | T | C | A | G | G | C | C | C | T | A | T | C | A | C | G |
| NA18870L | C | A | T | T | A | G | T | A | C | G | A | A | T | A | G | G | C | G | T | A | C | G | T | C | G | C | G | T | A | A | G | T | C | A | G | G | C | C | C | C | A | T | C | A | C | G |
| NA18924L | C | A | T | T | A | G | T | A | C | G | A | A | T | A | G | G | C | G | T | A | C | G | T | C | G | C | G | C | A | A | G | T | C | A | G | A | C | C | C | C | A | T | C | A | C | G |
| NA18940R | G | A | T | T | G | G | T | A | C | G | A | A | T | A | G | G | C | C | T | A | C | G | T | C | G | C | G | T | A | A | G | C | C | A | G | G | C | C | C | C | A | T | C | A | C | G |
| NA18962R | C | A | T | T | A | G | T | A | C | G | A | A | T | A | G | G | C | G | T | A | C | G | T | C | G | C | G | T | A | A | G | T | C | A | G | G | C | C | C | T | A | T | C | A | C | G |
| NA19347L | G | A | T | T | G | G | T | A | C | G | A | A | T | A | G | G | C | C | T | A | C | G | T | C | G | C | G | T | A | A | G | C | C | A | G | G | C | C | C | C | A | T | C | A | C | G |
| NA19372L | G | A | T | T | G | G | T | A | C | G | A | A | T | A | G | G | C | C | T | A | C | G | T | C | G | C | G | T | A | A | G | C | C | A | G | G | C | C | C | C | A | T | C | A | C | G |
| NA19383R | C | A | T | T | G | G | T | A | C | G | A | A | T | A | G | G | C | G | T | A | C | G | T | C | G | C | G | T | A | A | G | C | C | A | G | G | C | C | T | C | A | T | C | A | C | G |
| NA19397R | C | A | T | T | G | G | T | A | C | G | A | A | T | A | G | G | C | G | T | A | C | G | T | C | G | C | G | T | A | A | G | C | C | A | G | G | C | C | C | C | A | T | C | A | C | G |
| NA19435R | C | A | T | T | G | G | T | A | C | G | A | A | T | A | G | G | C | G | T | A | C | G | T | C | G | C | G | T | A | A | G | C | C | A | G | G | C | C | T | C | A | T | C | A | C | G |
| NA19467R | C | A | T | T | G | G | T | A | C | G | A | A | T | A | G | G | C | G | T | A | C | G | T | C | G | C | G | T | A | A | G | C | C | A | G | G | C | C | C | C | A | T | C | A | C | G |
| NA19704R | G | A | T | T | G | G | T | A | C | G | A | A | T | A | G | G | C | G | T | A | C | G | T | C | G | C | G | T | A | A | G | C | C | A | G | G | C | C | C | C | A | T | C | A | C | G |
| NA20336L | C | A | T | T | G | G | T | A | C | G | A | A | T | A | G | T | C | G | T | A | C | G | T | C | G | C | G | T | A | A | G | C | C | A | G | G | C | C | C | C | A | T | C | A | C | G |
| NA20336R | C | A | T | T | A | G | T | A | C | G | A | A | T | A | G | G | C | G | T | A | C | G | T | C | G | C | G | T | A | A | G | T | C | A | G | G | C | C | C | C | G | T | C | A | C | G |
| NA20509L | C | A | T | T | G | G | T | A | C | G | A | A | T | A | G | G | C | C | T | A | C | G | T | C | G | C | G | T | A | A | G | C | C | A | G | G | C | C | C | C | A | T | C | A | C | G |
| NA20582R | C | A | T | T | A | G | T | A | C | G | A | A | T | A | G | G | C | G | T | A | C | G | T | C | G | C | G | T | A | A | G | T | C | A | G | G | C | C | C | C | A | T | C | A | C | G |
| Denisova | G | A | T | T | G | G | T | A | C | G | A | A | T | A | G | G | C | G | T | A | C | G | T | C | G | C | G | T | A | A | G | C | C | A | G | G | C | C | C | C | A | T | C | A | C | G |
| AltaiNeandrtl | G | A | T | T | G | G | T | A | C | G | A | A | T | A | G | G | C | G | T | A | C | G | T | C | G | C | G | T | A | A | G | C | C | A | G | G | C | C | C | C | A | T | C | A | C | G |

Fig. s4

Xp11hs

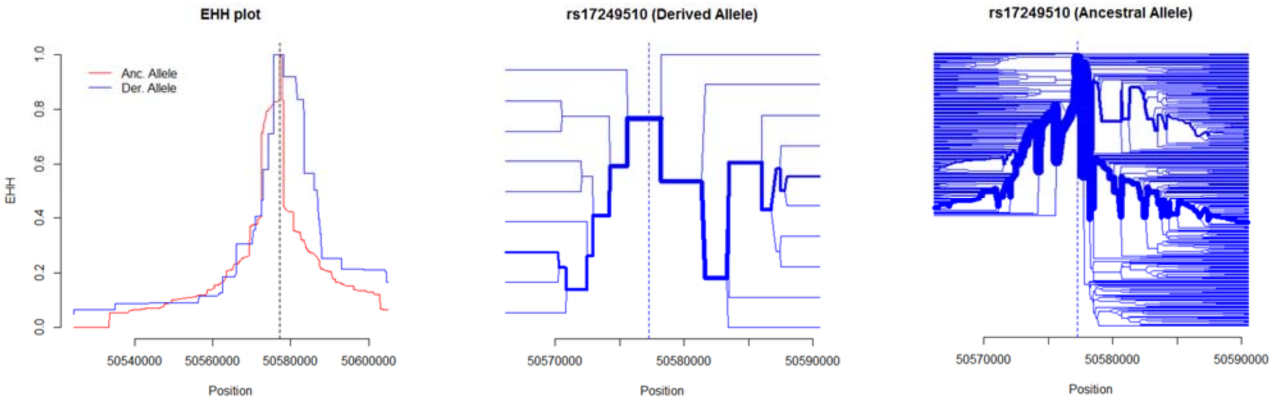


dys44

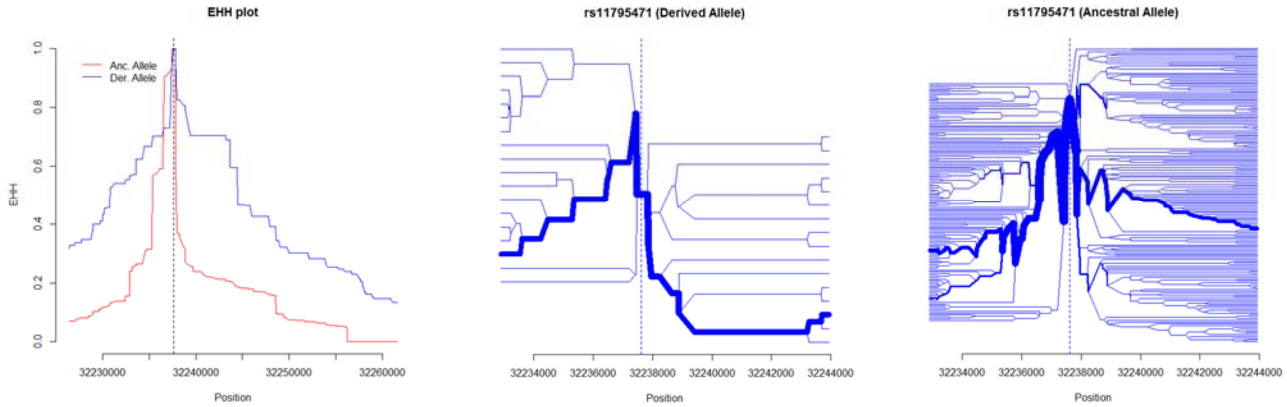

RRM2P4

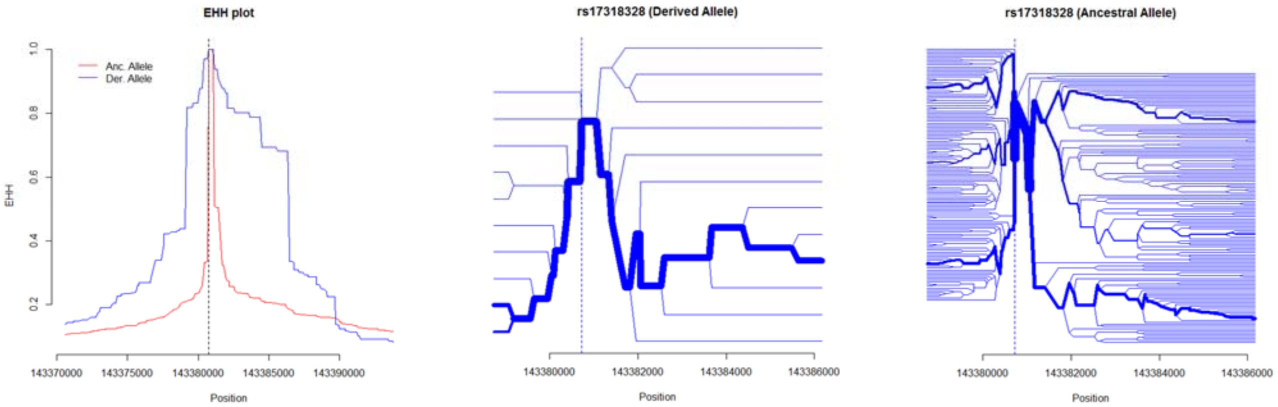


17q21inv

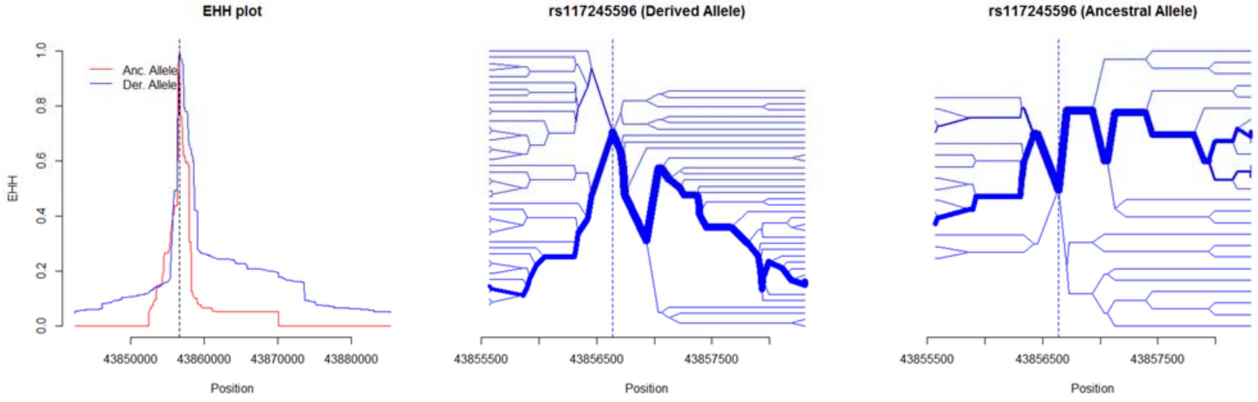

STAT2

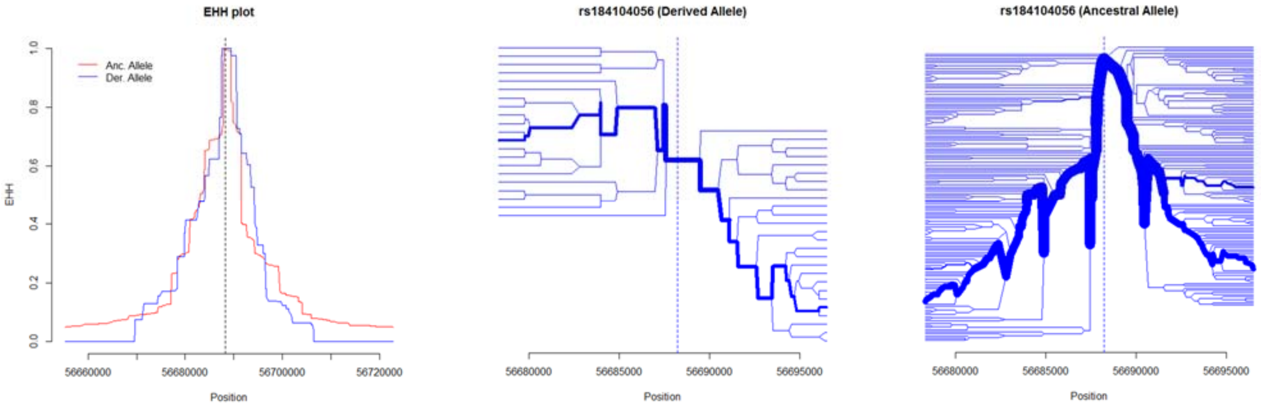
EHH plot
Anc. Allele
Der. Allele
EHH
0.0
0.2
0.4
0.6
0.8
1.0
56660000
56680000
56700000
56720000
Position
rs184104056 (Derived Allele)
56680000
56685000
56690000
56695000
Position
rs184104056 (Ancestral Allele)
56680000
56685000
56690000
56695000
Position

OAS

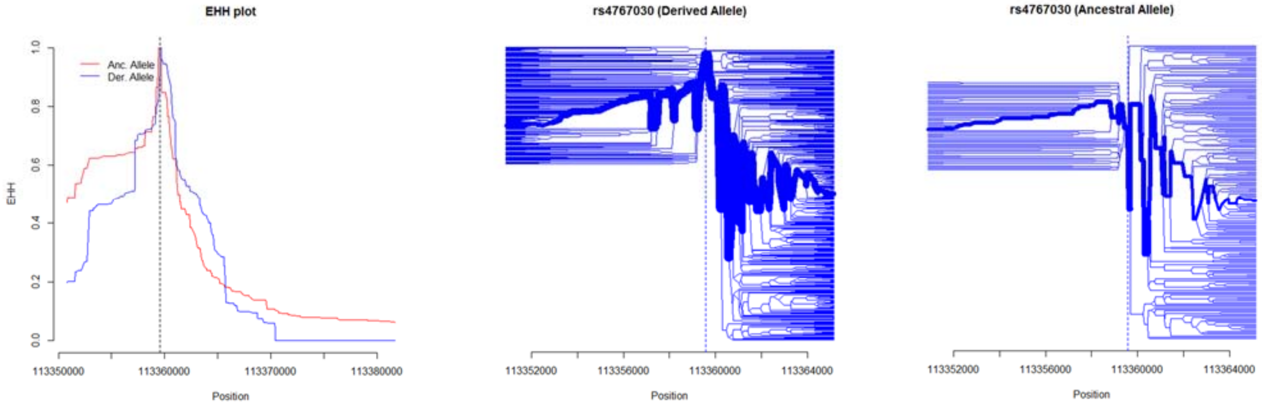
EHH plot
Anc. Allele
Der. Allele
EHH
0.0
0.2
0.4
0.6
0.8
1.0
113350000
113360000
113370000
113380000
Position
rs4767030 (Derived Allele)
113352000
113356000
113360000
113364000
Position
rs4767030 (Ancestral Allele)
113352000
113356000
113360000
113364000
Position

HYAL

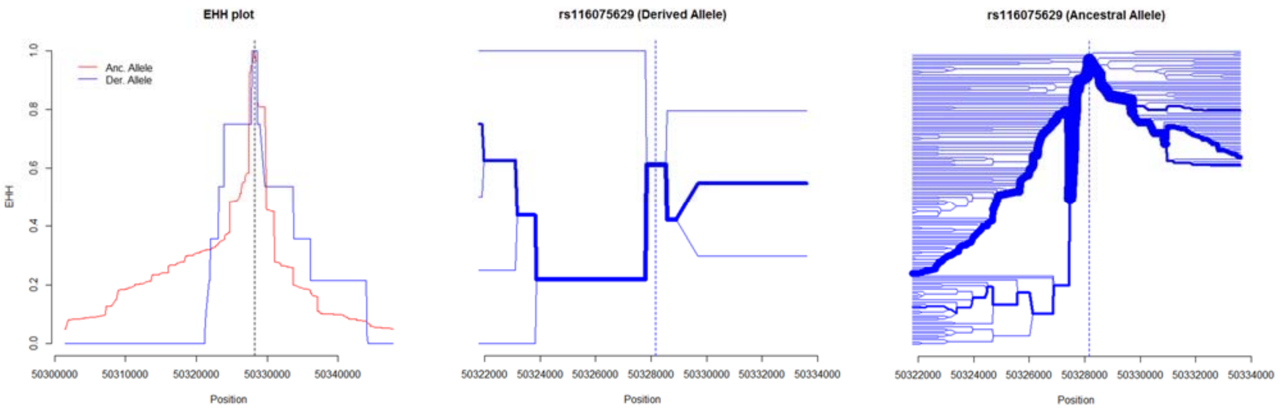
EHH plot
Anc. Allele
Der. Allele
EHH
Position
rs116075629 (Derived Allele)
Position
rs116075629 (Ancestral Allele)
Position

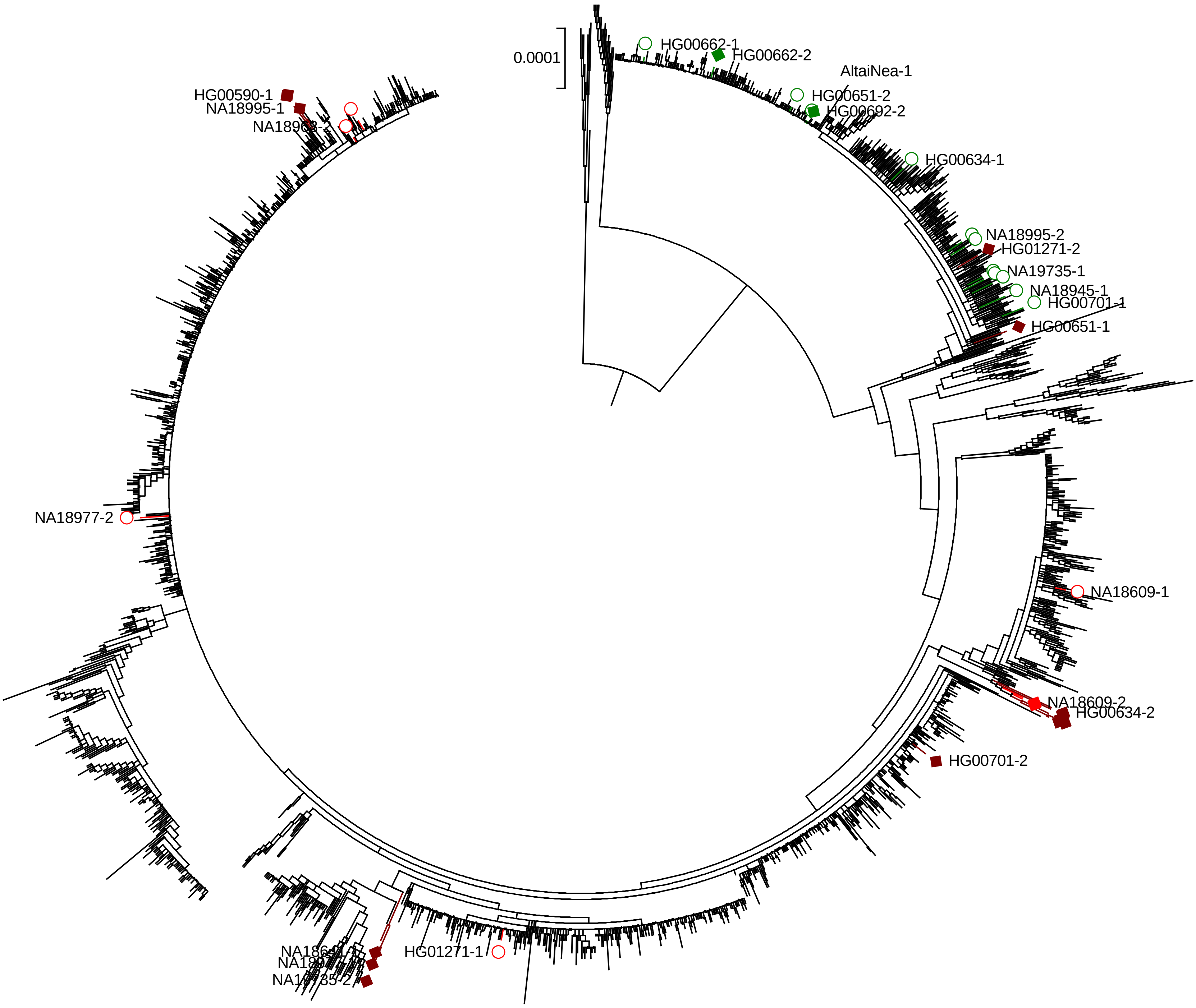
0.0001
HG00662-1
HG00662-2
AltaiNea-1
HG00651-2
HG00692-2
HG00634-1
NA18995-2
HG01271-2
NA19735-1
NA18945-1
HG00701-1
HG00651-1
NA18609-1
NA18609-2
HG00634-2
HG00701-2
HG01271-1
NA18977-2
HG00590-1
NA18995-1
